# The glia club: Validation of polarization biomarkers for human microglia (HMC3) using quantitative real time RT-qPCR

**DOI:** 10.1101/2024.09.26.615202

**Authors:** Esmé S. M. Franck, Jasmyne A. Storm, Jueqin Lu, Mon Francis M. Obtial, Sanoji Wijenayake

## Abstract

Microglia are the primary immune cells of the brain and play critical roles in neurodevelopment, neuroprotection, the maintenance of homeostasis, and neurotoxicity. Classification of microglia polarization, however, remains contentious. Identifying suitable biomarkers for gene expression analysis of microglia polarization is crucial for characterizing the biological significance of microglia in health and disease. In this study, we use human microglia clone 3 (HMC3) cells to validate and test suitable internal controls (*GAPDH*, *PKM*, *18S, ACTB, PGK1, TKT1, TPI1*), homeostatic (*CD68, TGF-β, IBA1, BIN1, RGS10*), proinflammatory (*IL-6, CXCL10, CCL5, SAA, IL-1β*) and anti-inflammatory (*CCL2, SOCS3, IL-10, CD200R1, ARG1*) markers along with their transcriptional profiles upon interferon-gamma (IFN-γ) stimulation to characterize the microglia polarization spectrum. Our study is the first to present a comprehensive list of biomarkers with detailed methodology on gene selection, primer design, RT-qPCR parameters, and transcript abundance in baseline and polarized HMC3 cells. Our study will increase the rigor of gene expression analysis and target selection in a widely used brain macrophage.

## Introduction

Microglia originate from progenitors in the embryonic yolk sac (Alliot et al., 1999) and form a self-renewing population in the adult brain (Butovsky & Weiner, 2018). After migrating to the developing central nervous system (CNS) during gastrulation, the amoeboid microglia shift to a ramified, homeostatic state (Ladeby et al., 2005). Microglia are the resident immune cells of the brain parenchyma and are highly responsive to environmental perturbations, stress, and stimuli (Perry et al., 1985). Microglia contribute to both innate and adaptive immune responses and are critical for maintaining a healthy CNS. They are primarily involved in brain development and neurogenesis, synaptic pruning, phagocytosis, autophagy and the secretion of cytokines, chemokines, growth factors, and reactive oxygen and nitrogen species to the surrounding environment (Lenz & Nelson, 2018). Thus, neuronal function is closely associated with microglia function. Upon stimulation, microglia become enlarged and acquire shortened processes irrespective of the type and duration of stress and/or stimuli (Ladeby et al., 2005). However, depending on the specificity of the stimuli, stress and duration, microglia may polarize into a “bushy”, rod-shaped, or hyper-ramified morphology and exert pro-inflammatory and/or anti-inflammatory effects in the CNS (Morrison et al., 2017; Walker et al., 2014). Therefore, microglia are one of the most dynamic and responsive macrophages in the CNS.

Polarized microglia produce exaggerated responses to inflammatory stimuli and immune challenges (Lima et al., 2022) and show changes in morphology, antigen presentation and proliferation (Perry & Holmes, 2014). To empirically study gene expression profiles of microglia, previous studies have used a variety of stimuli to prime cells (Harry & Kraft, 2008). One of the most common agents used to induce polarization is lipopolysaccharide (LPS), a bacterial endotoxin derived from *Salmonella typhimurium*. However, original experiments done on HMC3 cells found that LPS (10 µg/ml, 24 hours) may not induce sufficient microglia polarization as it induced little or no increase in *IL-6* titers (Janabi et al., 1995). Also, LPS induced polarization may be an outcome of intraventricular LPS injection that leads to autophagosome formation rather than a direct immune response (Ye et al., 2020). In terms of functional relevance, a site-specific injection of LPS to the brain is not very conducive to biological processes that polarize microglia in the CNS (Lively & Schlichter, 2018). Further, immortalized human and rodent microglia appear less responsive to LPS compared to primary cultures (Dello Russo et al., 2018). Therefore, alternates to LPS are often used in HMC3 polarization experiments. A leading alternative is the use of cytokines/chemokines, such as tumor necrosis factor-alpha (TNF-α) and IFN-γ (Benveniste & Benos, 1995).

TNF-α modulates the expression of class I and II major histocompatibility complex (MHC), increases cytokine expression (*e.g.* IL-6), and promotes transcription factor activity, including nuclear factor kappa-light-chain-enhancer of activated B cells (NF-κB), activator protein 1 (AP1), and interferon regulatory factor-1 (IRF-1) (Benveniste & Benos, 1995). However, HMC3 cells do not endogenously secrete TNF-α, even after stimulation with LPS or IL-1α (Janabi et al., 1995). However, HMC3 cells endogenously produce IFN-γ following LPS treatment (Shahbazi et al., 2018). In HMC3, IFN-γ treatment can increase the expression of MHCII, activation markers CD11b and CD68, and chemokine receptors CCR3 and CXCR3 (Dello Russo et al., 2018). Additionally, microglia exposed to IFN-γ have increased inducible nitric oxide synthase (iNOS) and IL-6 production (Kann et al., 2022). Therefore, we ultimately chose IFN-γ as the stimulant to induce microglia polarization in our study.

The HMC3 cell line was established from 8-10-week-old embryos transfected with the SV40 T antigen (Janabi et al., 1995). ATCC assessed morphology and karyotype, used PCR to confirm the human origin of the cells without other neuroglia (*e.g.* astrocytes), and distributes the immortalized cell line. Antigen expression and cytokine production in HMC3 are similar to that of the primary microglia cells prior to transfection (Janabi et al., 1995) and experiments performed by Dello Russo et al. (2018) confirmed that the HMC3 cell line retains most of its original antigenic profiles. As such, the HMC3 cell line is an ideal *in vitro* model that can be used to delineate the molecular underpinnings of human microglia polarization and the corresponding biological outcomes.

Numerous studies have described microglia polarization with “M1” and “M2” terminology, where “M1” represents a pro-inflammatory state and “M2” represents an anti-inflammatory state (Mantovani et al., 2004; Mills et al., 2000). However, given that the definition of microglia polarization itself is not well established, classifying microglia as “M1” or “M2” is over-simplistic (Ransohoff, 2016). In the current next generation sequencing era, neuroglia research has entered a dynamic phase, where single cell genomics, transcriptomics, and proteomics combined with high throughput imaging technologies have provided empirical insight into a dynamic and stimuli dependent polarization spectrum.

In this study, we present a comprehensive list of biomarkers that can be used to characterize homeostatic, pro-inflammatory, and anti-inflammatory states within the microglia polarization spectrum using the HMC3 cell line in response to 10 and 50 ng/ml IFN-γ treatment. We also tested seven candidate reference genes, including glyceraldehyde-3-phosphate dehydrogenase (*GAPDH)*, pyruvate kinase isozyme (*PKM*), 18S small-subunit ribosomal rRNA (*18S*), actin-beta (*ACTB*), phosphoglycerate kinase 1 (*PGK1*), transketolase 1 (*TKT1*), and triosephosphate isomerase 1 (*TPI1*). The homeostatic markers analyzed were cluster of differentiation 68 (*CD68*), transforming growth-factor beta (*TGF-β*), ionized calcium binding adaptor molecule 1 (*IBA-1*), bridging integrator-1 (*BIN1*), and regulator of G-protein signaling 10 (*RGS10*). The pro-inflammatory markers analyzed were interleukin-6 (*IL-6*), C-X-C motif chemokine ligand 10 (*CXCL10*), C-C motif chemokine ligand 5 (*CCL5*), serum amyloid A (*SAA*), and interleukin-1 beta (*IL-1β*). The anti-inflammatory markers analyzed were C-C motif chemokine ligand 2 (*CCL2*), suppressor of cytokine signaling 3 (*SOCS3*), interleukin-10 (*IL-10*), cluster of differentiation 200 receptor 1 (*CD200R1*), and arginase 1 (*ARG1*).

Our study found that *GAPDH, PKM,* and *18S* are suitable reference genes that can be used in IFN-γ treated human microglia for transcript normalization. A spectrum of homeostatic (*CD68, TGF-β*), pro-inflammatory (*IL-6, CXCL10, CCL5, SAA*), and anti-inflammatory (*CCL2, SOCS3*) biomarkers changed in transcript abundance in response to IFN-γ priming, emphasizing the phenotypic plasticity of microglia and the absence of concrete “M1” and “M2” states, where pro-inflammatory markers robustly increase and anti-inflammatory markers generally decrease in transcript abundance with polarization. Our study is the first to present detailed methodology for target selection, primer design, primer testing, and transcript abundance for a comprehensive selection of internal controls, homeostatic, pro-inflammatory, and anti-inflammatory biomarkers in response to IFN-γ treatment in the HMC3 cell line. Our study offers valuable insight for gene expression analysis and biomarker selection in the HMC3 cell line.

## Materials and Methods

### 1.0 Human microglia cell (HMC3) culture

One vial of Passage (P) 1 HMC3 at 2.26×10^6^ cells/ml (ATCC: CRL-3304; lot/batch number: 70026037) was donated by Dr. Patrick O. McGowan (University of Toronto) to Dr. Sanoji Wijenayake (The University of Winnipeg). The vial was delivered on dry ice on February 14, 2023. All required permits and material transfer agreements from ATCC and The Canadian Food Inspection Agency (compliance letter ID: CL-2022-0019-4) were obtained prior to the transfer. A certificate of analysis was provided by ATCC at the time of the original purchase certifying the absence of mycoplasma and cells from other species and tissue sources. The culture was tested for mycoplasma contamination using the MycoFluor Mycoplasma Detection Kit (ThermoFisher Scientific: M7006) as per manufacturer instructions prior to usage at The University of Winnipeg. P1 cells were immediately seeded upon delivery in a T25 culture flask with 10 ml of complete growth media consisting of Minimum Essential Medium (EMEM) (ATCC: 30-2003) and 10% Fetal Bovine Serum (FBS) (ATCC: 30-2020) and incubated at 37°C with 5% CO_2_ as per manufacturer recommendations. Complete media was replaced every 48 hours to avoid excessive acidification (Dello Russo et al., 2018). Cell confluence was determined qualitatively through observations by two independent operators using the EVOS XL Core imaging system (Invitrogen: AMEX1000). Once the cells reached >90% confluence, they were transferred to a T75 flask and maintained until reaching >90% confluence, approximately 4 days. Subsequently, cultures were sub-cultivated (1:3 ratio) to generate P2-P10 passages. All passages were cryopreserved in 10% DMSO (Bioshop: DMS666.100) and 90% complete media at −150°C. 0.25% Trypsin with 0.52 nM EDTA (ATCC: 30-2101) solution was used for cell detachment and harvesting.

### 2.0 IFN-γ treatments for microglia polarization

The pro-inflammatory, type II, T-lymphocyte cytokine, IFN-γ, was used to prime HMC3 cells. P8 cells were seeded and cultured in T75 flasks and upon reaching >90% confluence, were plated onto 6-well culture plates at 2×10^5^ cells/well (concentration determined using the Countess 3 automated cell counter; Invitrogen: AMQAX2000) for downstream experiments (n=5 biological replicates/treatment). Once cells reached >90% confluence, a subset of cells were treated with either 10 or 50 ng/ml IFN-γ (Millipore-Sigma: SRP3058) diluted in complete media, with untreated HMC3 cells as a baseline control. Cells were incubated for 24 hours at 37°C and 5% CO_2_ post-treatment, harvested and stored at −80°C for downstream analyses (**Figure 1**).

**Fig. 1.**
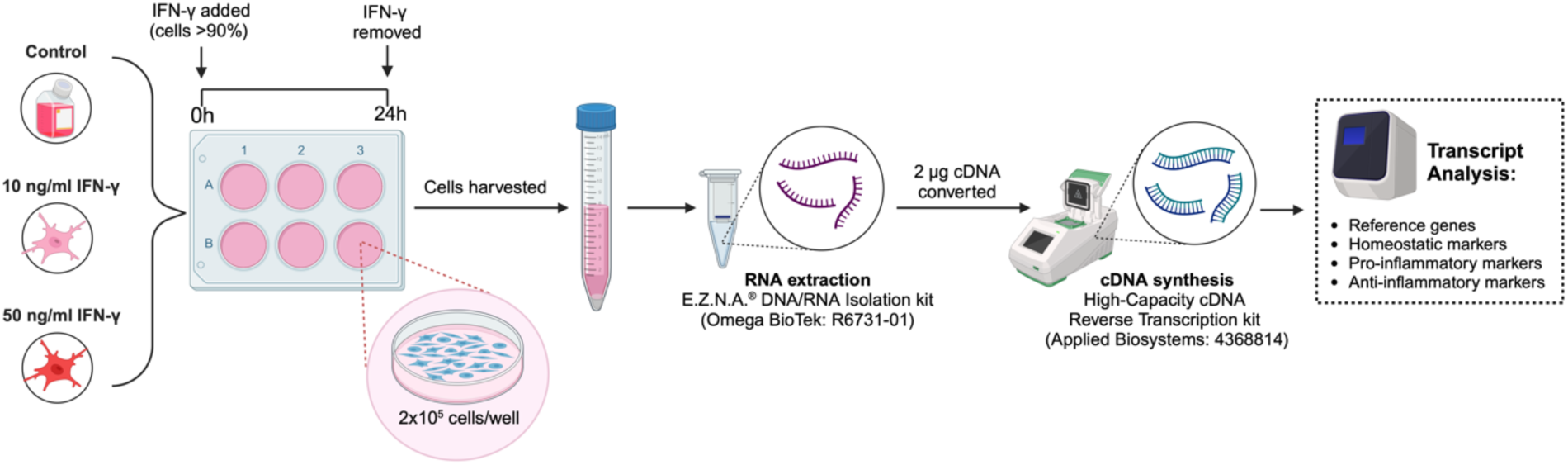
Schematic of human culturing and IFN-γ treatment. HMC3 cells were plated at 2×10^5^ cells/well. At >90% confluence, a subset of cells was treated with 10 ng/ml IFN-γ or 50 ng/ml IFN-γ. Untreated cells are used as controls. 24 hours post-IFN-γ treatment, all cells were harvested, total soluble RNA was extracted, and 2 μg were reverse transcribed into cDNA. RT-qPCR was used to analyze transcript abundance of reference genes, homeostatic, pro-inflammatory, and anti-inflammatory microglial markers.

### 3.0 Cell viability and mitochondrial output

#### 3.1 Brightfield microscopy

To assess morphology, HMC3 cells were imaged via brightfield microscopy prior to harvesting. Images were obtained using the EVOS XL Core imaging system with the 10x and 20x objective lenses.

#### 3.2 Live and dead cell counting

10 μl aliquots of P8 HMC3 cells were diluted (1:1, v/v) with 0.4% trypan blue stain (Invitrogen: T10282). The cell suspension was loaded onto a reusable glass slide (Invitrogen: A25750) and total cell concentration, live cell concentration/percentage, and dead cell concentration/percentage were determined using the Countess 3 automated cell counter under default settings.

##### 3.3 3 -[4,5-dimethylthiazol-2-yl]-2,5 diphenyl tetrazolium bromide (MTT) assay

A CyQUANT MTT cell viability assay (Invitrogen: V-13154) was used to quantify mitochondrial output of HMC3 cells with and without IFN-γ treatment as per manufacturer instructions. Absorbances were read at 570 nm using a BioTek Synergy H1 Multimode microplate reader (Agilent: BTSH1M2SI). Prior to quantification, a standard curve was used to determine the optimal cell density. 1×10^3^ cells/well, 1×10^4^ cells/well, 5×10^4^ cells/well and 1×10^5^ cells/well (n=3 biological replicates/treatment) were tested. Cells were treated with 10 or 50 ng/ml IFN-γ for 24 hours, with untreated HMC3 cells used as a baseline control, as previously described. Based on the results from the pilot study, 2.5×10^4^ cells/well was used for subsequent MTT cell viability assays (n=5 biological replicates/treatment).

### 4.0 RNA extractions and quantifications

Total soluble RNA (≥18 nucleotides) was extracted from frozen cell pellets using the E.Z.N.A.^®^ DNA/RNA Isolation kit (Omega BioTek: R6731-01) according to manufacturer instructions. RNA concentration (ng/µl) and purity (A260:280 and A260:230) were measured using a NanoDrop One/One^C^ Microvolume-UV/Vis spectrophotometer (ThermoFisher Scientific: ND-ONE-W). RNA purity was determined by the A260:280 and A260:230. Samples with ratios of 1.8-2.0 were used for complementary DNA (cDNA) synthesis. RNA Clean & Concentrator kit (Zymo Research: R1017) was used to clean samples with A260:280 <1.8-2.0 according to manufacturer instructions. RNA stability and integrity were verified by resolving samples on a 1% TAE-agarose gel (1:1, v/v, 2x RNA loading dye) (Life Technologies: R0641) stained with Red Safe dye (Froggabio: 21141) using a Sub-Cell GT Agarose Gel Electrophoresis System (BioRad: 1704401) **(Supplementary Figure S1)**.

### 5.0 cDNA synthesis

2000 ng of total soluble RNA was reverse transcribed using the High-Capacity cDNA Reverse Transcription kit (Applied Biosystems: 4368814) according to manufacturer instructions. A T100™ Thermal Cycler (BioRad: 1861096) was used for synthesis using the following amplification parameters: 25°C for 10 mins; 37°C for 120 mins; 85°C for 5 mins; hold at 4°C.

### 6.0 Primer design

Primers were purchased from Qiagen or Eurofins Genomics. Primers were designed using nucleotide sequence information available at the National Center for Biotechnology information (NCBI): http://www.ncbi.nlm.nih.gov or obtained from previously published research. NCBI-designed primers were selected using Primer-BLAST against the coding region of the reference sequence from the generated options. Primer pairs were assessed for compliance with the Minimum Information for Publication of Quantitative Real-Time PCR Experiments (MIQE) guidelines (Bustin et al., 2009) using sequence information available from NCBI and the OligoAnalyzer^™^ Tool from Integrated DNA Technologies (IDT, Coralville, Iowa, USA).

The following parameters were used to test for compliance: 20-25 nucleotides in length, melting temperatures between 55-65°C, a difference in melting temperature <3°C between forward and reverse primers, hairpin temperature below 45°C, GC content between 40-60%, self-dimerization ΔG< 15%, and heterodimerization ΔG< 20% (Bustin et al., 2009). To ensure efficient amplification by the AmpliTaq^®^ Fast DNA polymerase, primer pairs were only selected if the intended product size was <350 nucleotides. Primer parameters are described in **Table 1**.

**Table 1.**
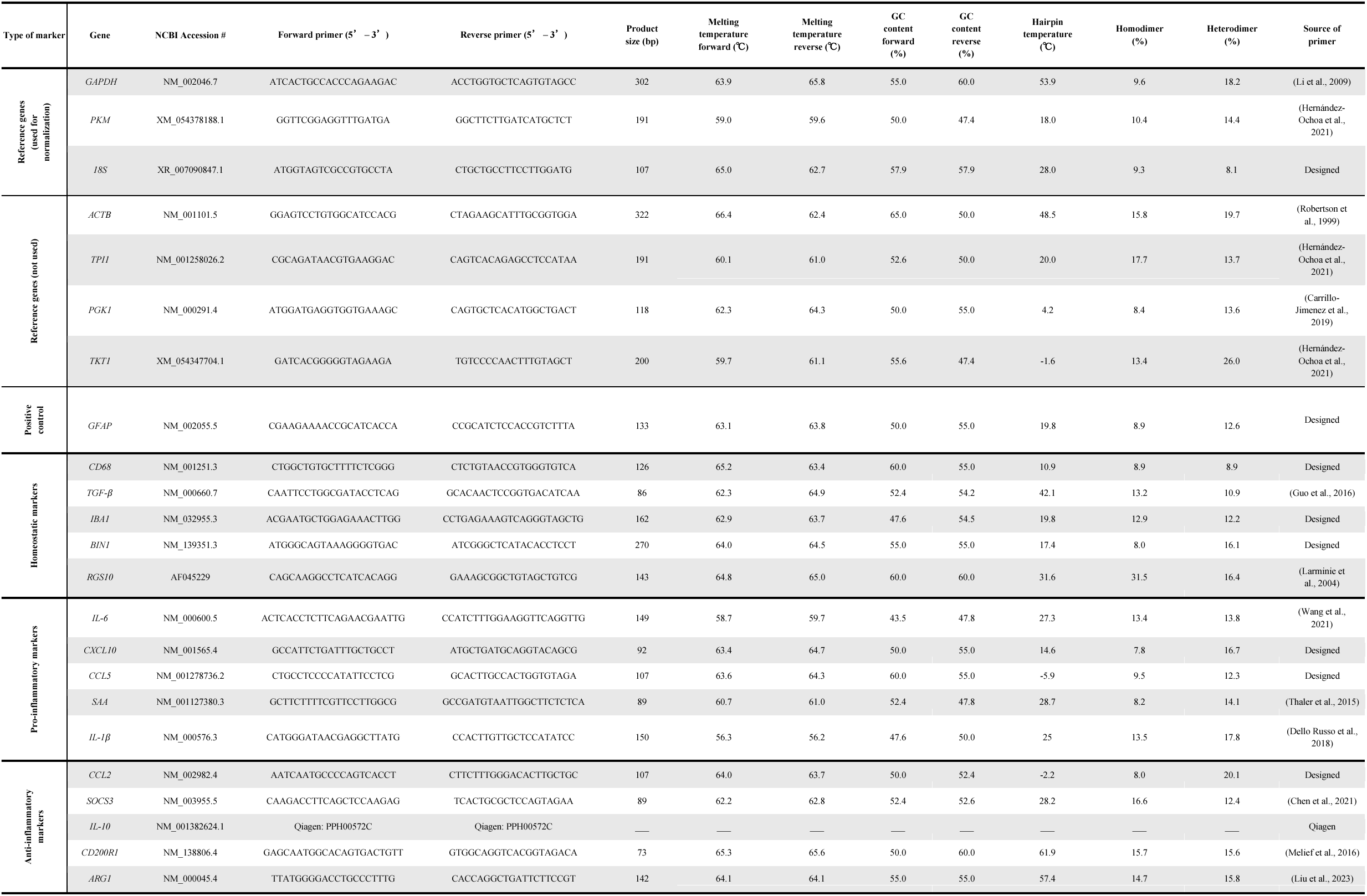
Primer parameters for RT-qPCR. Targets are identified by gene name, NCBI accession number, targeted transcript variants, and forward and reverse primer sequences. Product size (bp), forward and reverse melting temperatures (°C) and GC content (%), hairpin temperature (°C), and homodimer and heterodimer values (%) are listed. The sources of the primers (designed using NCBI or obtained from literature) are indicated. All primers target *Homo sapiens*.

### 7.0 Quantitative reverse transcription polymerase chain reaction (RT-qPCR)

RT-qPCR was performed using the QuantStudio^™^ 5 PCR system (Applied Biosystems: A28139; 96-well, 0.2 ml block) with Fast SYBR^™^ Green PCR chemistry (Applied Biosystems: 4385612). Annealing temperatures for primer pairs were determined by testing a range of temperatures approximately ± 5°C of the predicted melting temperature of the forward primer (5’-3’), using the Veriflex setting of the QuantStudio^™^ 5 PCR system. Temperature testing was conducted using a pool sample (10 ng/µl) that represents all test samples. Melt curve analysis was conducted to determine the specificity of the primer pairs. Primer pairs that generated a single, sharp melt peak void of primer dimers and a derivative reporter >200,000 were used for quantification. An 8-point standard curve ranging from 500 ng/μl to 3.91 ng/μl was used to determine the optimal cDNA amount. Analyses were done in triplicate and an inter-plate converter was used to account for variability between plates and runs.

Seven reference genes were tested: *GAPDH, PKM, 18S*, *ACTB*, *PGK1*, *TKT1*, and *TPI1*. *GAPDH, PKM*, and *18S* have the highest thermal stability and lowest intra/intergroup variability in response to IFN-γ treatment, as determined by NormFinder software (Andersen et al., 2004) (RRID: SCR_003387). All targets were normalized against the geometric mean of the three reference genes. The astrocyte marker, glial fibrillary acidic protein (*GFAP*), was quantified to determine specificity of the cells towards microglia and to illustrate the absence of astroglia in cultures. *Rattus norvegicus* hypothalamic cDNA (30 ng/μl) was used as a positive control to illustrate that the lack of amplification reported in HMC3 samples was not due to non-functioning primers. The primers used for *GFAP* quantification are 100% conserved between human and rat. Relative transcript levels are denoted as mean with individual points, n=5 biological replicates per experimental condition and three technical replicates per biological replicate. Transcript abundance of candidate homeostatic (*CD68, TGF-β, IBA-1, BIN1, RGS10*), pro-inflammatory (*IL-6, CXCL10, CCL5, SAA, IL-1β*), and anti-inflammatory (*CCL2, SOCS3, IL-10, CD200R1, ARG1*) biomarkers were analyzed using the comparative Ct method (2^- ΔΔCt^).

The RT-qPCR parameters for primers that were not used for quantification in the study yet remain potential options are represented in **Supplementary Table S1** and **Supplementary Table S2**, respectively. Ultimately, these primers were not selected for quantification if a second primer pair for the same target amplified earlier, produced higher percent efficiency, had sharper melt peaks, or illustrated a higher derivative reporter during temperature and standard curve testing. A complete list of unspecific primers that failed to amplify the targets are listed in **Supplementary Table S3**.

### 8.0 Statistical analysis

Statistical analysis was done using SPSS (IBM; version 29.0.2.0), and figures were created using Prism (GraphPad; version 9.5.1), BioRender.com, and RStudio (Posit; version 2023.06.1+524). Normality testing was conducted using Shapiro-Wilk tests, as the sample size is less than 30. Levene’s tests were used to assess homogeneity of variance. All data are normally distributed (p > 0.05) and parametric. As such, a one-way analysis of variance (ANOVA) was used to determine significant differences across groups with 95% confidence intervals (p < 0.05). Extreme outliers with an interquartile range >3 was identified using the SPSS boxplot outlier function and removed from the dataset, when necessary. No more than one outlier was removed per treatment group. Pairwise comparisons between groups were determined using Tukey HSD post-hoc tests (p < 0.05). Raw RT-qPCR and MTT assay data are available in **Supplementary Table S4**.

## Results

### 1.0 HMC3 morphology, cell viability and live/dead cell counts in response to IFN-γ

Morphological changes of microglia have been reported upon stimulation with specific ligands and cytokines/chemokines, where chronic polarization typically results in shorter branches and amoeboid-shaped, enlarged cell bodies, whereas ramified, homeostatic microglia tend to exhibit complex networks and a highly branched morphology (H. Wang et al., 2023). However, 10 and 50 ng/ml IFN-γ treatment leads to minimal changes in morphology, when compared to control cells (**Figure 2a**). Specifically, we observe a slight increase in amoeboid-shaped microglia with IFN-γ treatment (particularly at the 50 ng/ml dosage), while most of the cells are ramified with network formation and branching. These connections are present in all treatment conditions.

**Fig. 2.**
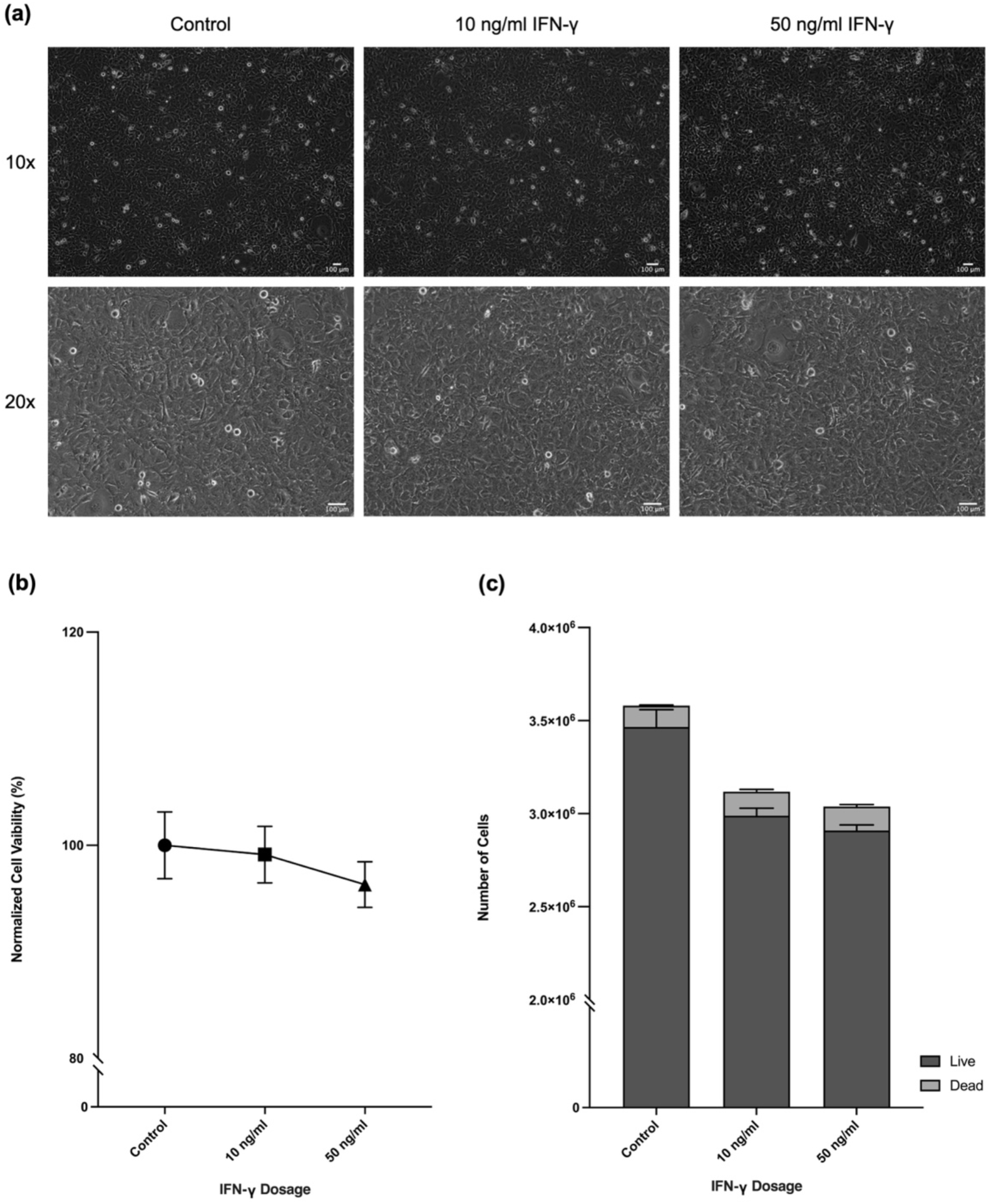
HMC3 cell viability with IFN-γ treatment. **(a)** Cell morphology determined via brightfield microscopy (EVOS XL Core Imaging System) at 10x and 20x. **(b)** MTT assay in control and IFN-γ treated cells. **(c)** Total number of live and dead cells in control and IFN-γ-treated groups.

We determined cell viability in response to IFN-γ using a commercial MTT assay. Cell viability remains unchanged between the control, 10 ng/ml IFN-γ, and 50 ng/ml IFN-γ cells (**Figure 2b**). Specifically, the percent cell viability (mean ± SEM) is 100% ± 3.12% for control cells, 99.13% ± 2.65% for the 10 ng/ml IFN-γ cells, and 96.32% ± 2.14% for the 50 ng/ml IFN-γ cells. We also determined the percentage of live and dead cells in response to IFN-γ using an automated cell counter to determine if the lack of changes in metabolic output measured via the MTT assay is a true representation of cell viability. Live and dead cell percentages (mean ± SEM) are 96.80% ± 0.37% live and 3.2% ± 0.37% dead for the control cells, 96.00% ± 0.89% live and 4.00% ± 0.89% dead for the 10 ng/ml IFN-γ cells, and 95.80% ± 1.16% live and 4.20% ± 1.16% dead for the 50 ng/ml IFN-γ cells. Total cell count (*F*_(2,12)_ = 2.531, p = 0.121), live cell count (*F*_(2,12)_ = 2.681, p = 0.109), and dead cell count (*F*_(2,12)_ = 0.081, p = 0.923) remain unchanged in response to 10 and 50 ng/ml IFN-γ treatments (**Figure 2c**).

### 2.0 HMC3 reference genes

We tested seven candidate genes, previously identified to be stable internal controls in neuroglia cells, in response to IFN-γ treatment, to determine the most suitable set of internal controls for normalization. The glycolytic enzyme, *GAPDH,* amplifies a single product (302 bp) at 63°C with 10 ng of cDNA input. Control cells amplify in the Ct range of 19.0-19.8. *PKM*, a second glycolytic enzyme, amplifies a single product (191 bp) at 63°C with 10 ng of cDNA input. Control cells amplify in the Ct range of 20.9-21.9. Finally, *18S*, which is involved in protein synthesis, amplifies a single product (107 bp) at 60°C with 10 ng of cDNA input. Control cells amplify in the Ct range of 10.7-12.3 (**Table 2**).

**Table 2.**
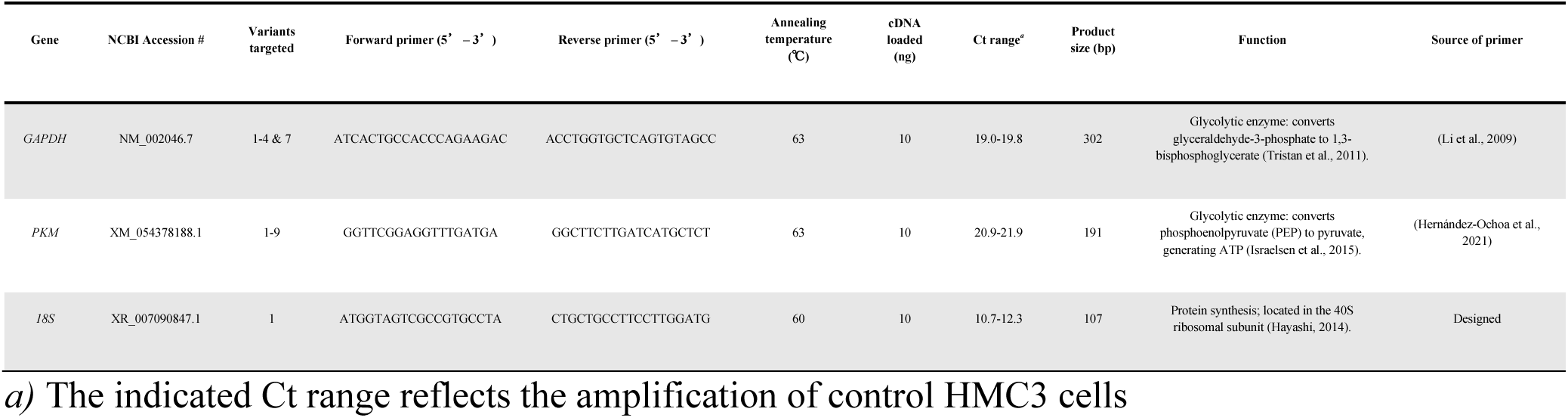
Primer sequences and RT-qPCR parameters for reference genes. Targets are identified by gene name, NCBI accession number, targeted transcript variants, forward and reverse primer sequences, annealing temperature (°C), cDNA amount (ng), threshold cycle (Ct), product size (bp), and function. The sources of the primers (designed using NCBI or obtained from literature) are indicated. All primers target *Homo sapiens*.

We determined the annealing temperature by testing which temperature had the highest transcript abundance with a single, sharp melt peak in the absence of primer dimers. The candidate reference genes in order of lowest to highest transcript abundance are: *PKM > GAPDH > 18S*. Further, *GAPDH* (*F*_(2,12)_ = 2.418, p = 0.131), *PKM* (*F*_(2,12)_ = 3.531, p = 0.062), and *18S* (*F*_(2,12)_ = 0.402, p = 0.678) remain unchanged in response to 10 and 50 ng/ml IFN-γ treatment (**Figure 3a**). However, the four additional reference candidates (**Supplementary Table S5**) significantly change in transcript abundance in response to IFN-γ treatment. Specifically, the cell cycle-regulator, *ACTΒ* (*F*_(2,11)_ = 7.416, p < 0.01) at 50 ng/ml IFN-γ (Tukey HSD, p < 0.01); the glycolytic enzyme, *PGK1* (*F*_(2,11)_ = 7.187, p = 0.010) at 10 ng/ml IFN-γ (Tukey HSD, p < 0.01); the glycolytic and gluconeogenic enzyme, *TKT1* (*F*_(2,12)_ = 11.994, p = 0.001) at 10 ng/ml IFN-γ (Tukey HSD, p < 0.01) and 50 ng/ml IFN-γ (Tukey HSD, p < 0.01); and the pentose phosphate pathway enzyme, *TPI1* (*F*_(2,11)_ = 13.280, p = 0.001) at 10 ng/ml IFN-γ (Tukey HSD, p = 0.001) and 50 ng/ml IFN-γ (Tukey HSD, p = 0.011) (**Figure 3b**). As such, we determined these to be unsuitable internal controls for RT-qPCR normalization (**Supplementary Figure S2**).

**Fig. 3.**
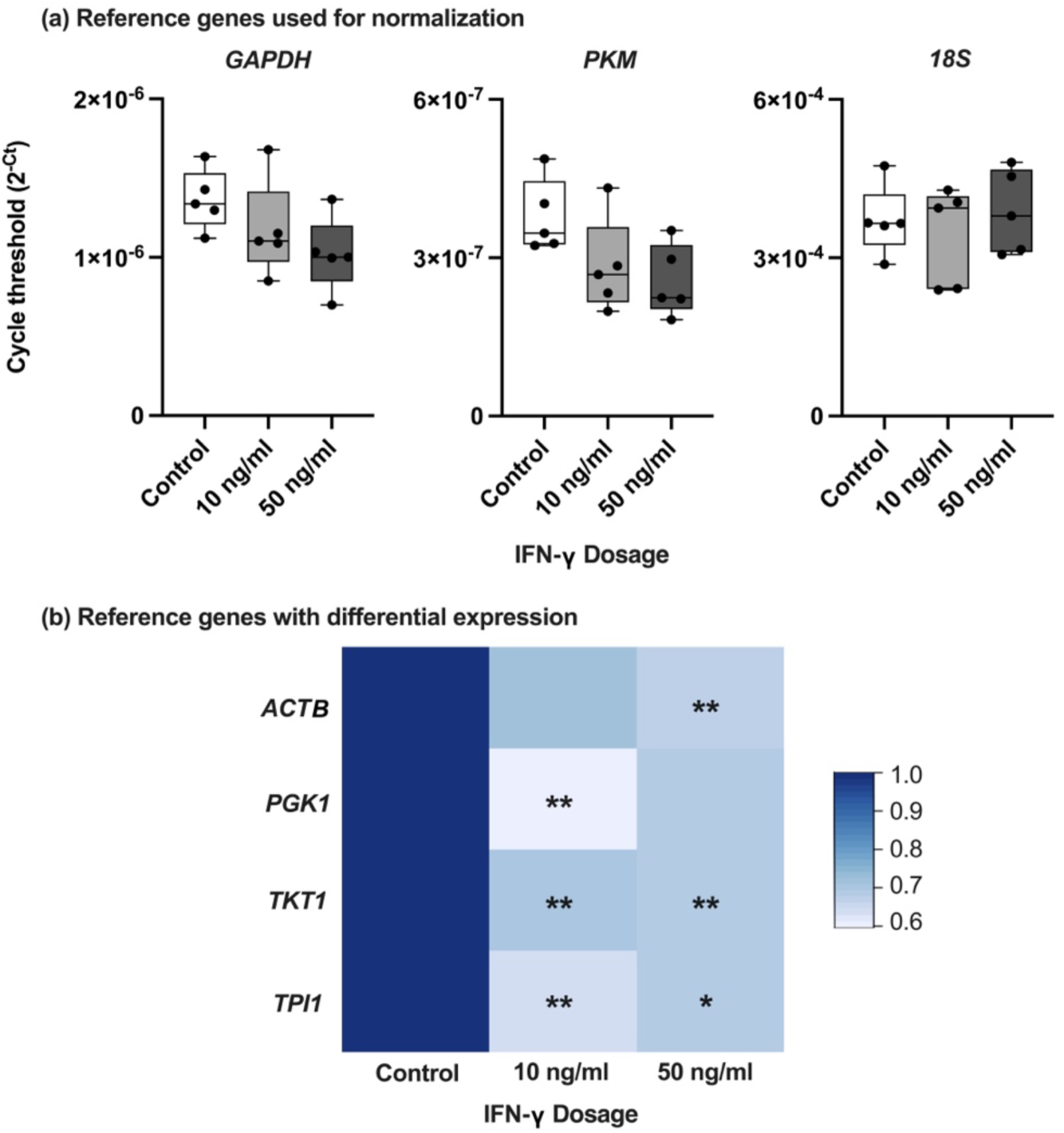
Relative transcript abundance of reference genes in HMC3 cells as determined by RT-qPCR in response to IFN-γ treatment. **(a)** Reference genes used for normalization (*GAPDH, PKM, 18S*). Data are mean 2^−Ct^ with individual points (n=5 biological replicates per experimental group). **(b)** Heat map of references genes with differential expression in responseto IFN-γ treatments (10 ng/ml and 50 ng/ml) compared to untreated controls. The 2^−Ct^values for each group (control, 10 ng/ml IFN-γ, and 50 ng/ml IFN-γ) were averaged (n=5 biological replicates per experimental group). Transcript abundance is represented as a fold change. Significant differences between treatment groups were determined using a one-way ANOVA with Tukey HSD (* p<0.05; ** p<0.01; *** p<0.001).

NormFinder ranks candidate reference genes according to their thermal stability and intragroup and intergroup variation (Andersen et al., 2004). According to NormFinder, *GAPDH* is the most stable gene, (thermal stability = 0.076), followed by *PKM* (thermal stability = 0.108) and *18S* (thermal stability = 0.155). The best combination of genes is *PKM* and *18S* (**Table 3**). For normalization of the target genes, we used the geometric mean of *GAPDH, PKM*, and *18S*.

**Table 3.** NormFinder results for *GAPDH*, *PKM*, and *18S* indicating thermal stability, intergroup variation, and intragroup variation. Best single gene and the best combination of two genes is indicated.

| Gene | Thermal stability value | Intragroup variation |  |  | Intergroup variation |  |  | Best gene | Best combination |
| --- | --- | --- | --- | --- | --- | --- | --- | --- | --- |
| *GAPDH* | 0.076 | 0.003 | 0.009 | 0.001 | 0.018 | 0.045 | -0.063 | ✓ | —— |
| *PKM* | 0.108 | 0.011 | 0.002 | 0.011 | 0.099 | -0.018 | -0.081 | —— | ✓ |
| *18S* | 0.155 | 0.001 | 0.047 | 0.023 | -0.118 | -0.027 | 0.144 | —— | ✓ |

### 3.0 HMC3 target genes

We quantified the transcript abundance of four homeostatic markers (*CD68, TGF-β, IBA1, BIN1, RGS10*), five pro-inflammatory markers (*IL-6, CXCL10, CCL5, SAA, IL-1β*), and six anti-inflammatory markers (*CCL2, SOCS3, IL-10, CD200R1, ARG1*). The relative mRNA abundance of each marker in the control, 10 ng/ml IFN-γ, and 50 ng/ml IFN-γ treated-cells is represented as a fold change relative to that of the control (**Figure 4a**) and the Ct distribution of all markers in the three conditions is illustrated (**Figure 4b**).

**Fig. 4.**
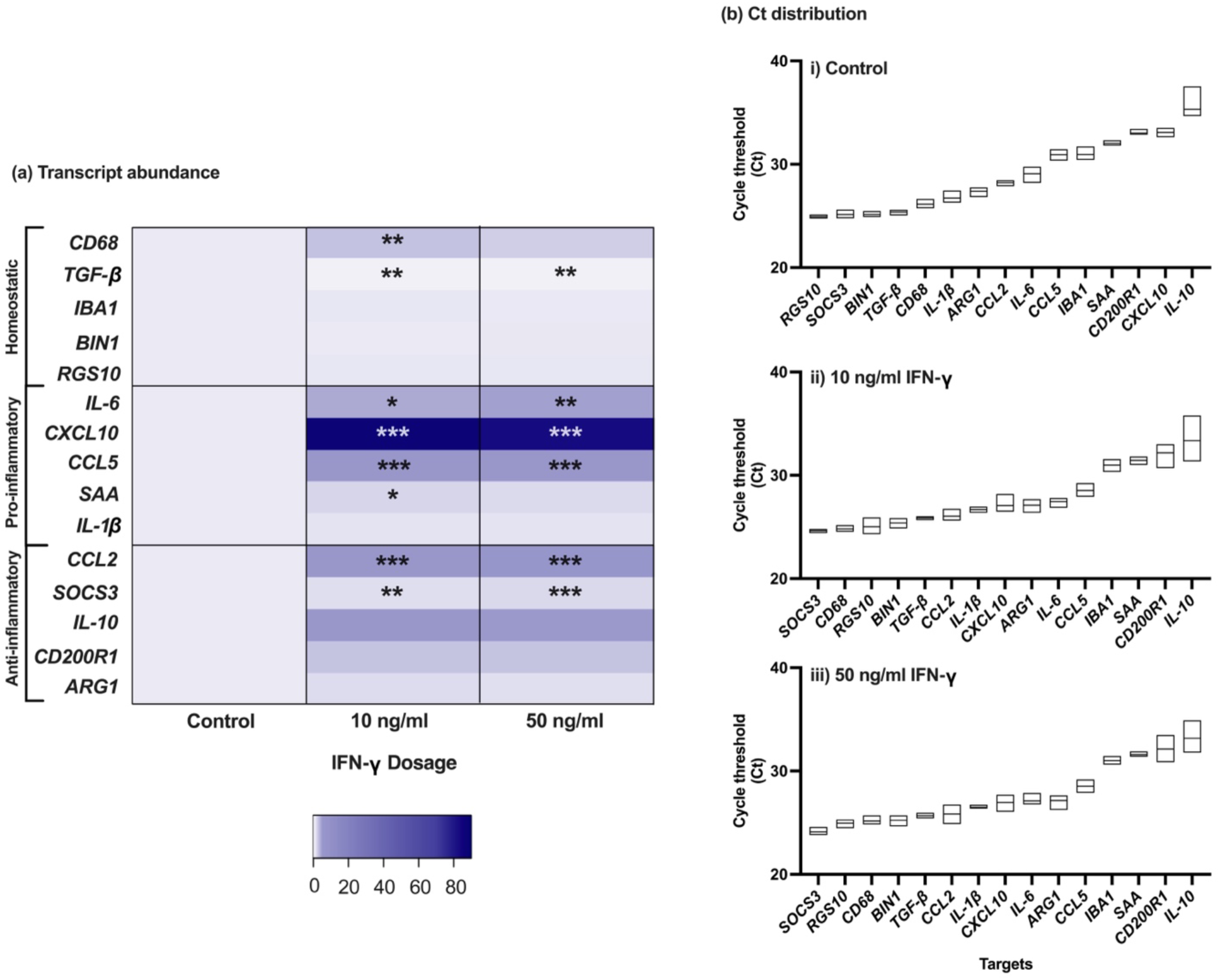
Expression profiles of homeostatic, pro-inflammatory, and anti-inflammatory biomarkers in HMC3 cells. **(a)**Transcript abundance of homeostatic markers (*CD68, TGF-β, IBA1, BIN1, RGS108*), pro-inflammatory markers (*IL-6, CXCL10, CCL5, SAA*, *IL-1β*) and anti-inflammatory markers (*CCL2, SOCS3, IL-10, CD200R1*, *ARG1*). The 2^−ΔΔC*t*^ values for each group (control, 10 ng/ml IFN-γ, and 50 ng/ml IFN-γ) were averaged (n=5 biological replicates per experimental group). Transcript abundance is represented as a fold change. Significant differences between treatment groups were determined using a one-way ANOVA with Tukey HSD (* p<0.05; ** p<0.01; *** p<0.001). **(b)** Threshold cycle (Ct) distribution under (i) Control, (ii) 10 ng/ml IFN-γ, and (iii) 50 ng/ml IFN-γ treated groups. Data are mean Ct with distribution (n=5 biological replicates per experimental group).

#### 3.1 Homeostatic markers

To identify microglia in their dynamic surveying state, we measured the transcript abundance of five markers previously identified to be expressed in homeostatic microglia. *CD68,* a transmembrane protein highly expressed in monocytes, amplifies a single product (126 bp) at 58°C with 20 ng of cDNA input. Control cells amplify in the Ct range of 25.9-26.8. The cytokine growth factor *TGF-β* amplifies a single product (86 bp) at 60°C with 50 ng of cDNA input. Control cells amplify in the Ct range of 25.0-25.6. The calcium-binding protein *IBA1* amplifies a single product (162 bp) at 60°C with 150 ng of cDNA input. Control cells amplified in the Ct range of 30.4-31.7. *BIN1*, which encodes for nucleocytoplasmic adaptor proteins, amplifies a single product (270 bp) at 64°C with 30 ng of cDNA input Control cells amplify in the Ct range of 24.6-25.5. Finally, *RGS10,* a regulator of G protein-coupled receptor signaling cascades, amplifies a single product (191 bp) at 62°C with 30 ng of cDNA input. Control cells amplify in the Ct range of 24.7-25.2 (**Table 4**).

**Table 4.**
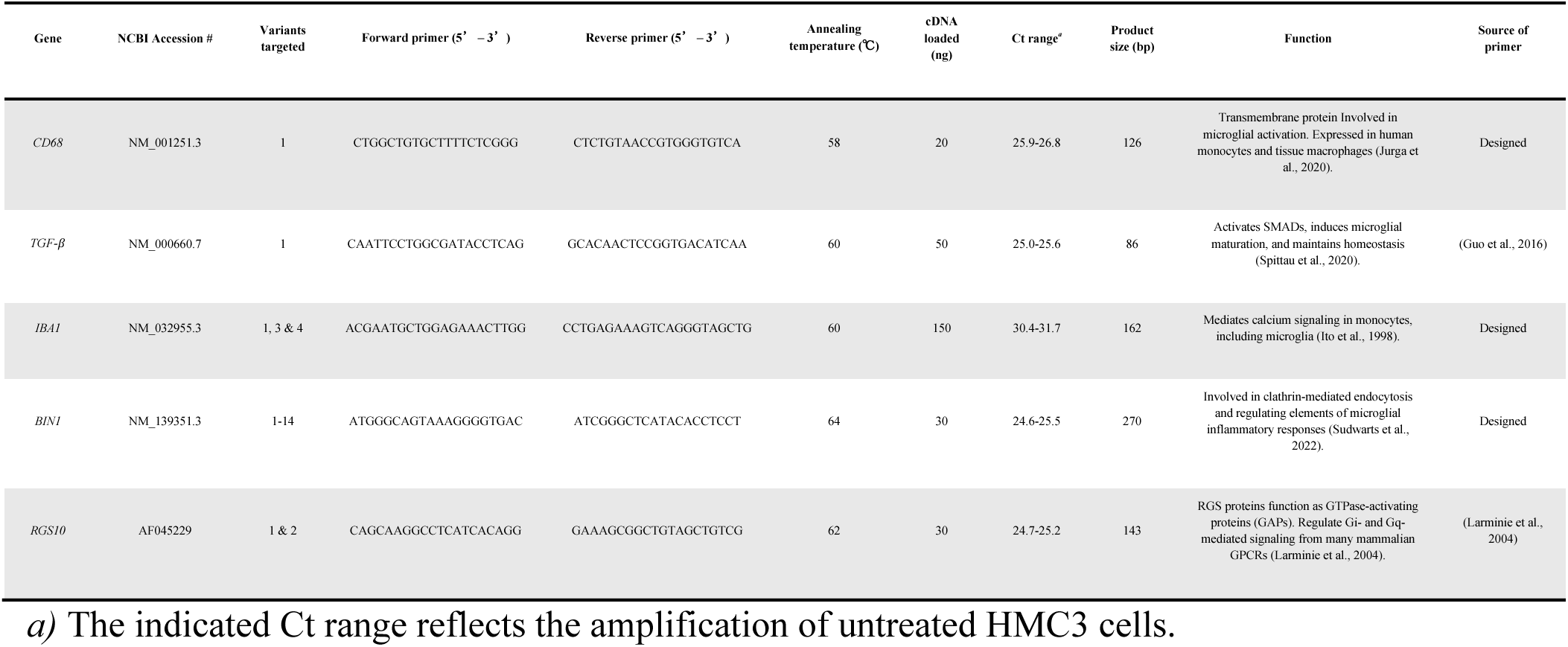
Primer sequences and RT-qPCR parameters for homeostatic genes. Targets are identified by gene name, NCBI accession number, targeted transcript variants, forward and reverse primer sequences, annealing temperature (°C), cDNA amount (ng), threshold cycle (Ct), product size (bp), and function. The sources of the primers (designed using NCBI or obtained from literature) are indicated. All primers target *Homo sapiens*.

*CD68*, increases with 10 ng/ml of IFN-γ treatment when compared to control cells (*F*_(2, 11)_ = 8.156, p < 0.01, Tukey HSD, p < 0.01), and remains unchanged between 10 and 50 ng/ml IFN-γ dosages (Tukey HSD, p = 0.360). Additionally, *TGF-β*, decreases in transcript abundance with IFN-γ (*F*_(2,12)_ = 10.027, p < 0.01) at 10 ng/ml (Tukey HSD, p < 0.01) and 50 ng/ml (Tukey HSD, p < 0.01), but remains unchanged between the two dosages (Tukey HSD, p = 0.881). The remaining homeostatic markers do not change with 10 or 50 ng/ml IFN-γ treatment: *IBA1* (*F*_(2,11)_ = 0.088, p = 0.917), *BIN1* (*F*_(2,12)_ = 1.467, p = 0.269), and *RGS10* (*F*_(2,12)_ = 1.088, p = 0.368) (**Figure 5a-e**).

**Fig. 5.**
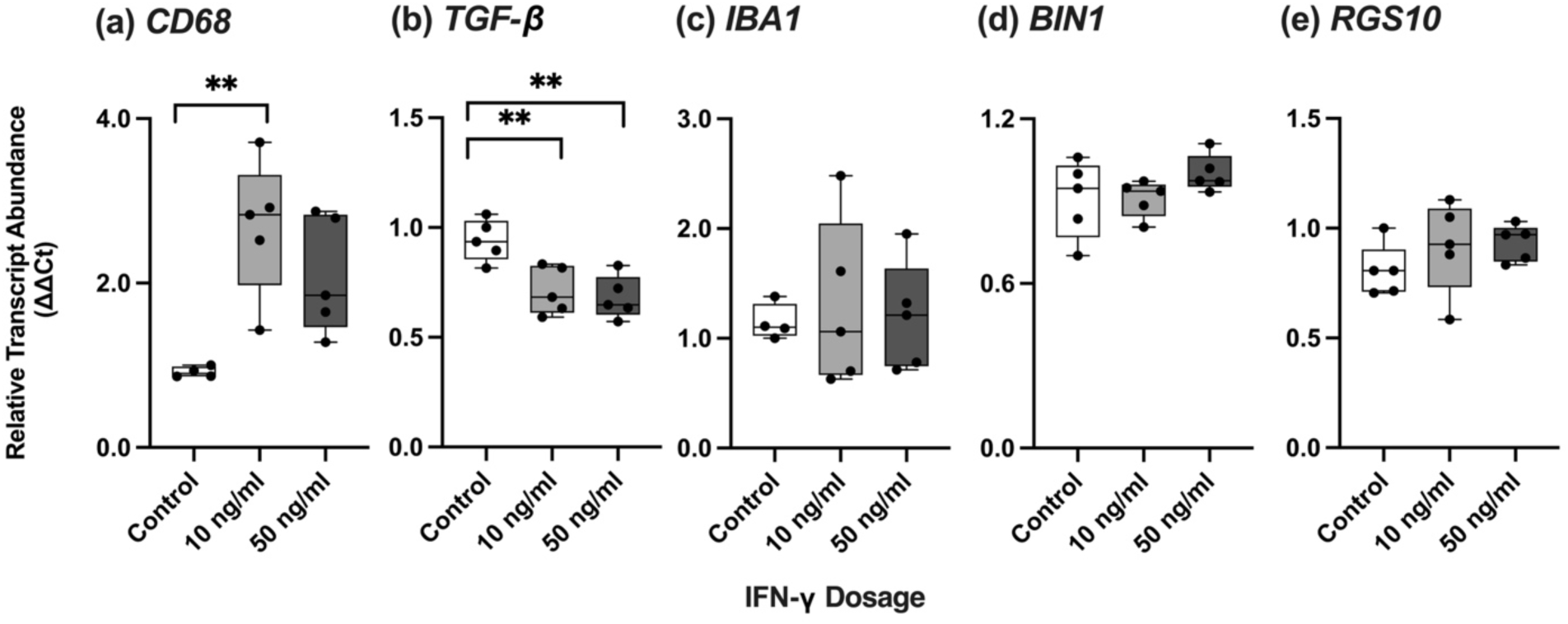
Relative transcript abundance of homeostatic markers in response to IFN-γ (10 ng/ml and 50 ng/ml) in HMC3 cells. **(a)** *CD68*, **(b)** *TGF-β*, **(c)** *IBA-1*, **(d)** *BIN1*, and **(e)** *RGS10*. Data are mean 2^−ΔΔC*t*^ with individual points (n=5 biological replicates per experimental group). Targets are normalized to the geometric mean of reference genes: *GAPDH*, *PKM* and *18S*. Significant differences between treatment groups were determined using a one-way ANOVA with Tukey HSD (* p<0.05; ** p<0.01; *** p<0.001).

We quantified the transcript abundance of an astroglia marker, *GFAP*, to test for cell-specificity. *GFAP* does not amplify in samples containing HMC3 cDNA inputs of 500 ng to 3.91 ng. The amplification profiles of the respective wells resemble those of the negative controls. However, the positive control containing *Rattus norvegicus* whole hypothalamic cDNA amplifies with an average Ct of 24.20 ± 0.18 (**Figure 6a-b**).

**Fig. 6.**
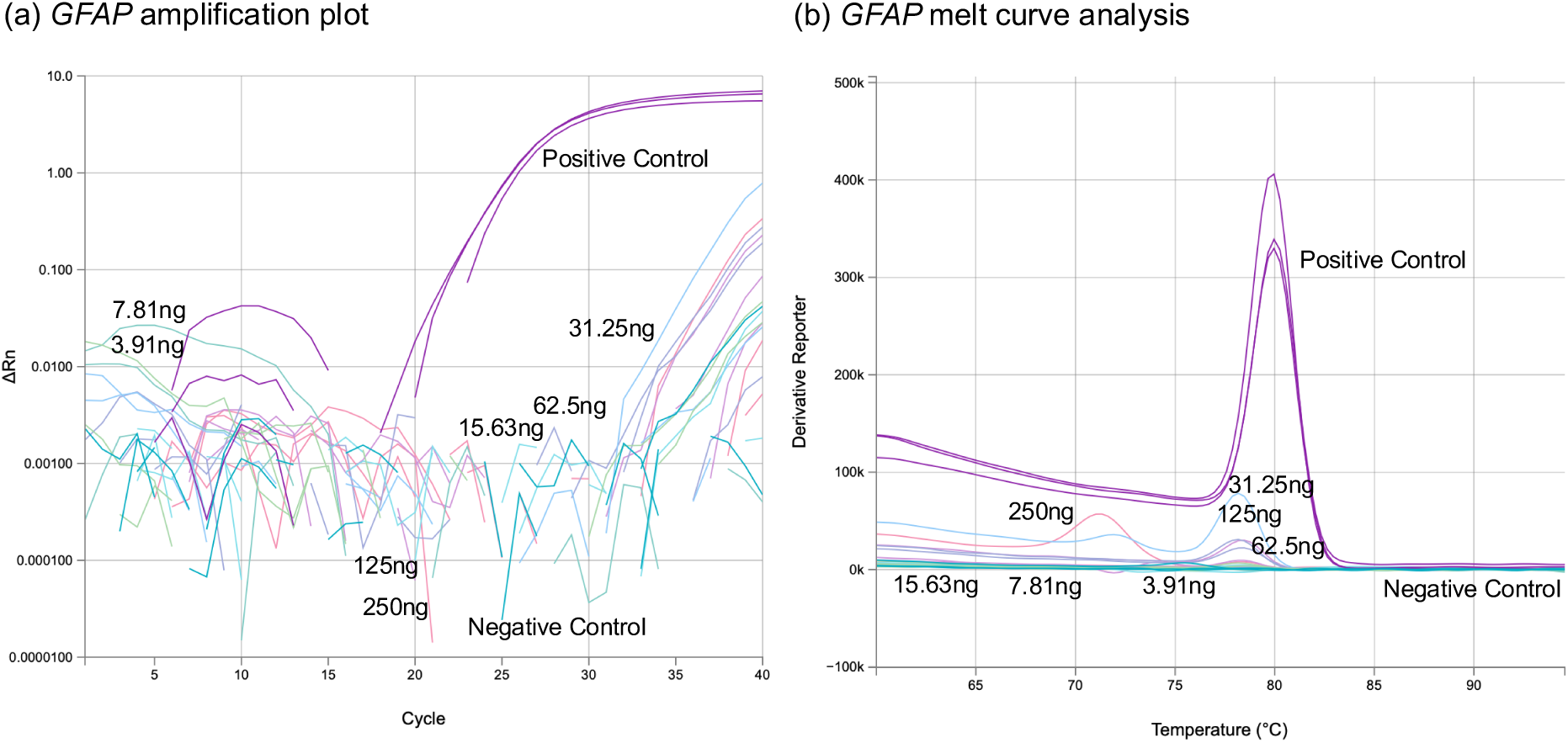
Expression profiles of the astrocyte marker, *GFAP*, in HMC3 cells. **(a)** Amplification plot and **(b)** melt curve analysis. Lack of amplification of the standards and negative control confirm the absence of astrocyte presence in HMC3. *Rattus norvegicus* hypothalamus cDNA was used as a positive control and illustrates amplification.

#### 3.2 Pro-inflammatory markers

Microglia can polarize to a pro-inflammatory state and regulate the expression of candidate genes. We measured the transcript abundance of genes previously known to be expressed by pro-inflammatory microglia in HMC3 cells (Jurga et al., 2020).

*IL-6,* a cytokine with pro- and anti-inflammatory properties, amplifies a single product (149 bp) at 60°C with 30 ng of cDNA input. Control cells amplify in the Ct range of 28.1-30.2. The pro-inflammatory chemokine *CXCL10* amplifies a single product (92 bp) at 59°C with 30 ng cDNA input. Control cells amplify in the Ct range of 32.2-33.8. The pro-inflammatory chemokine *CCL5* amplifies a single product (107 bp) at 63°C with 30 ng of cDNA input. Control cells amplify in the Ct range of 30.2-31.7. The apolipoprotein *SAA* amplifies a single product (89 bp) at 56°C with 50 ng of cDNA input. Control cells amplify in the Ct range of 32.3-33.8. Finally, the pro-inflammatory cytokine *IL-1β* amplifies a single product (150 bp) at 62°C with 50 ng cDNA input. Control cells amplify in the Ct range of 26.3-27.6 (**Table 5**).

**Table 5.**
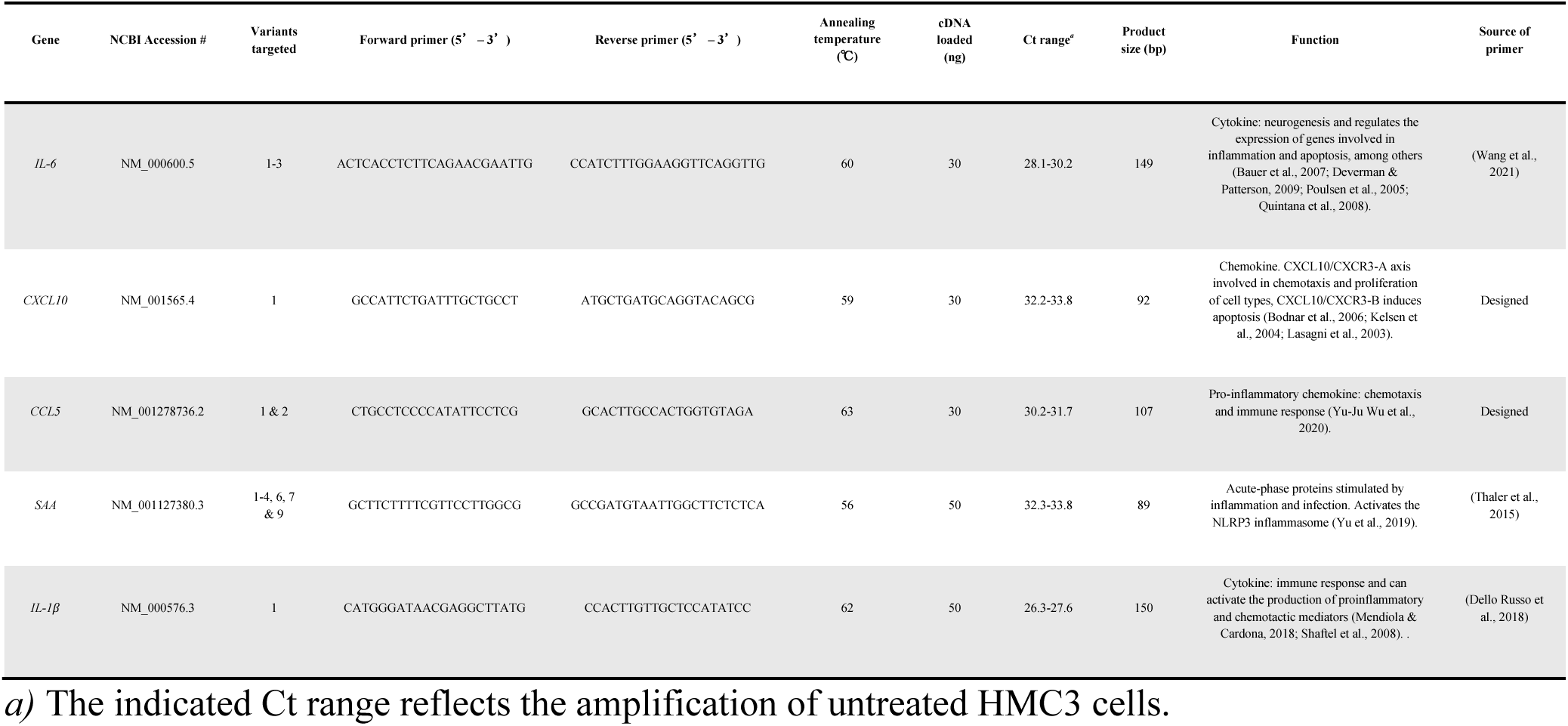
Primer sequences and RT-qPCR parameters for pro-inflammatory genes. Targets are identified by gene name, NCBI accession number, targeted transcript variants, forward and reverse primer sequences, annealing temperature (°C), cDNA amount (ng), threshold cycle (Ct), product size (bp), and function. The sources of the primers (designed using NCBI or obtained from literature) are indicated. All primers target *Homo sapiens*.

*IL-6* increases in transcript abundance with IFN-γ when compared to control cells (*F*_(2, 11)_ = 10.318, p < 0.01) at 10 ng/ml (Tukey HSD, p = 0.014) and 50 ng/ml (Tukey HSD, p < 0.01), but remains unchanged between the two dosages (Tukey HSD, p = 0.857). *CXCL10* also increases in abundance with IFN-γ treatment (*F*_(2,11)_ = 60.384, p < 0.001) at 10 ng/ml (Tukey HSD, p < 0.001) and 50 ng/ml (Tukey HSD, p < 0.001), but remains unchanged between the two dosages (Tukey HSD, p = 0.893). *CCL5* increases in abundance with IFN-γ when compared to control cells (*F*_(2,10)_ = 105.357, p < 0.001) at 10 ng/ml (Tukey HSD, p < 0.001) and 50 ng/ml (Tukey HSD, p < 0.001), but remains unchanged between the two dosages (Tukey HSD, p = 0.940). *SAA* increases in abundance with IFN-γ when compared to control cells (*F*_(2,11)_ = 6.459, p = 0.014) at 10 ng/ml (Tukey HSD, p = 0.012), but remains unchanged in cells that received the 50 ng/ml treatment when compared to the controls (Tukey HSD, p = 0.065). *SAA* transcript abundance remains unchanged between the two IFN-γ dosages (Tukey HSD, p = 0.567). *IL-1β* remains unchanged in response to both 10 and 50 ng/ml IFN-γ treatments (*F*_(2,11)_ = 0.806, p = 0.471) (**Figure 7a-e**).

**Fig. 7.**
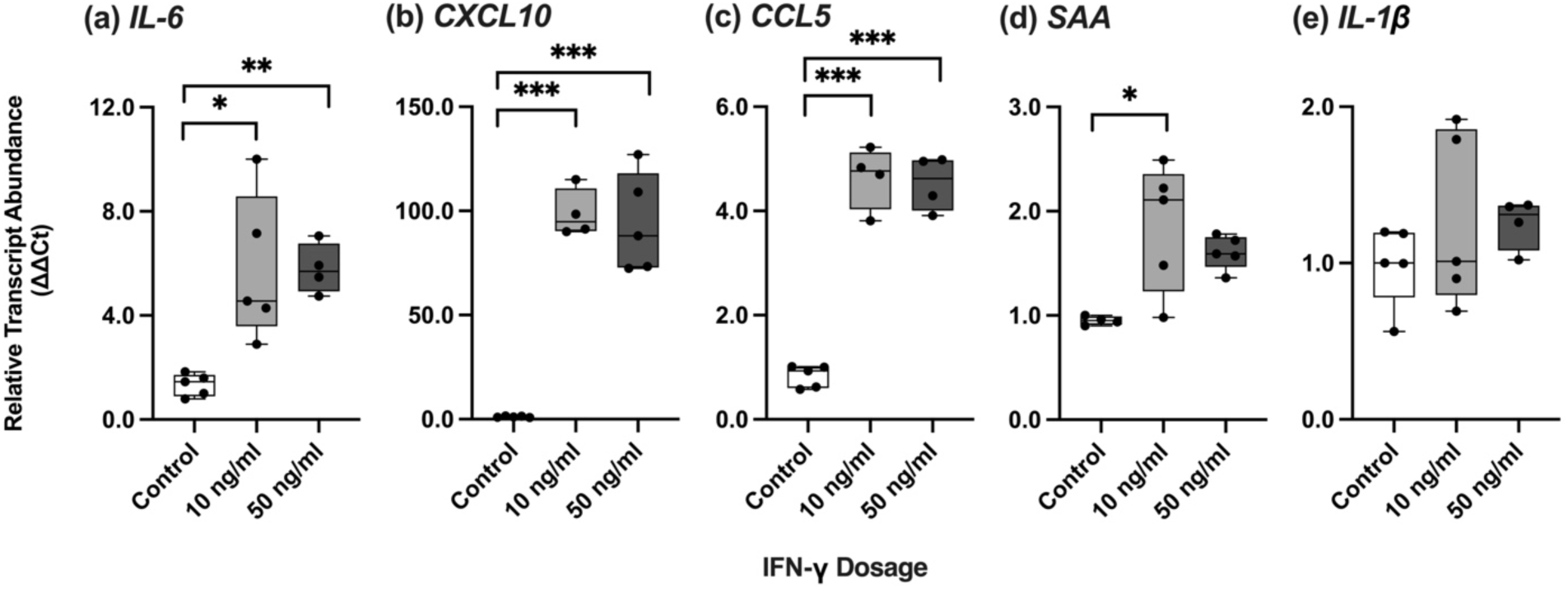
Relative transcript abundance of pro-inflammatory markers in response to IFN-γ (10 ng/ml and 50 ng/ml) in HMC3 cells. **(a)** *IL-6*, **(b)** *CXCL10*, **(c)** *CCL5*, **(d)** *SAA*, and **(e)** *IL-1β*. Data are mean 2^−ΔΔC*t*^ with individual points (n=5 biological replicates per experimental group). Targets are normalized to the geometric mean of reference genes: *GAPDH, PKM* and *18S*. Significant differences between treatment groups were determined using a one-way ANOVA with Tukey HSD (* p<0.05; ** p<0.01; *** p<0.001).

#### 3.3 Anti-inflammatory markers

We also measured the transcript abundance of genes previously known to be expressed by anti-inflammatory microglia in HMC3 cells. The anti-inflammatory chemokine *CCL2* amplifies a single product (107 bp) at 60°C with 30 ng cDNA input. Control cells amplify in the Ct range of 27.8-28.8. *SOCS3,* a negative regulator of cytokine signaling, amplifies a single product (89 bp) at 60°C with 50 ng of cDNA input. Control cells amplify in the Ct range of 24.7-25.6. The cytokine *IL-10* amplifies a single product at 60°C with 100 ng of cDNA input. Control cells amplify in the Ct range of 33.6-39.6. *CD200R1*, a receptor involved in inhibiting the pro-inflammatory phenotype of microglia, amplifies a single product (73 bp) at 60°C with 75 ng cDNA input. Control cells amplify in the Ct range of 32.3-34.0. Finally, *ARG1,* a regulator of nitric oxide levels, amplifies a single product (142 bp) at 64°C with 100 ng cDNA input. Control cells amplify in the Ct range of 26.6-28.0 (**Table 6**).

**Table 6.**
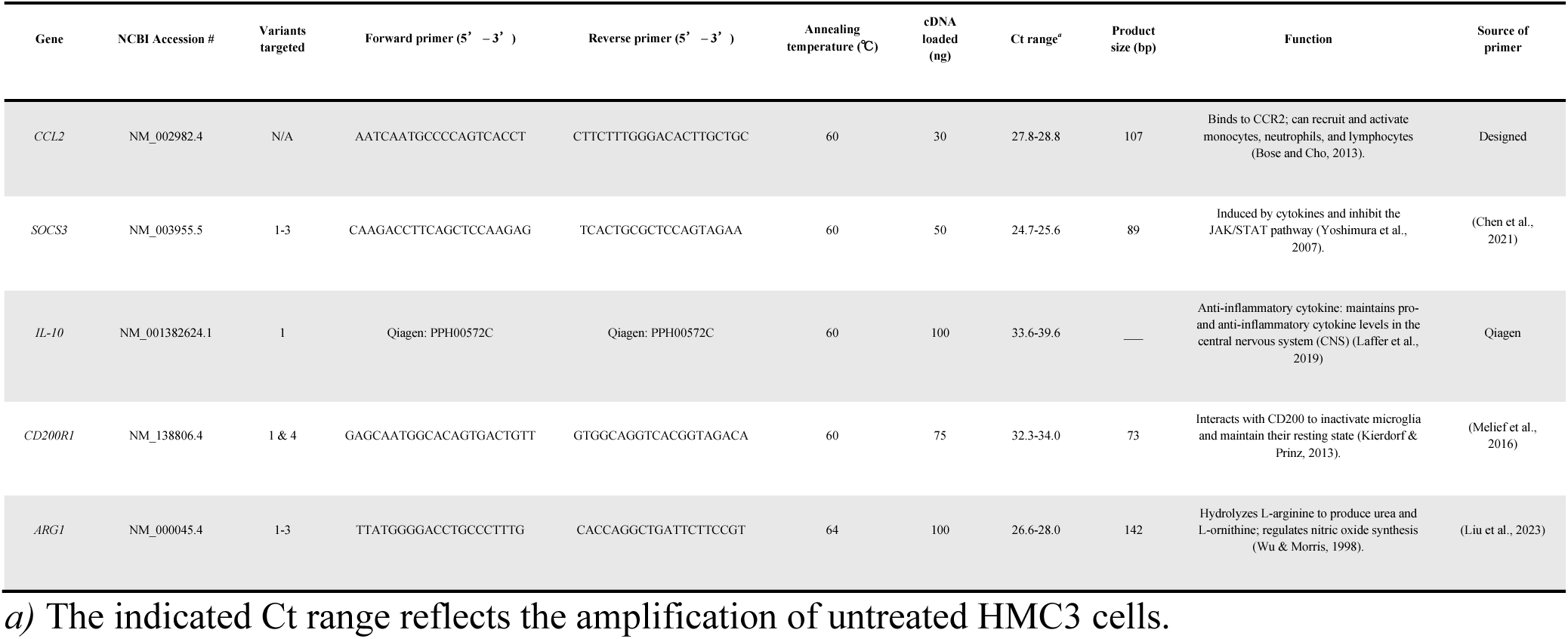
Primer sequences and RT-qPCR parameters for anti-inflammatory genes. Targets are identified by gene name, NCBI accession number, targeted transcript variants, forward and reverse primer sequences, annealing temperature (°C), cDNA amount (ng), threshold cycle (Ct), product size (bp), and function. The sources of the primers (designed using NCBI or obtained from literature) are indicated. All primers target *Homo sapiens*.

In response to IFN-γ treatment, two of five candidate anti-inflammatory markers change in transcript abundance in response to IFN-γ, whereas three remain unchanged. *CCL2* increases in transcript abundance with IFN-γ when compared to control cells (*F*_(2,12)_ = 27.659, p < 0.001) at 10 ng/ml (Tukey HSD, p < 0.001) and 50 ng/ml (Tukey HSD, p < 0.001), but remains unchanged between the two dosages (Tukey HSD, p = 0.474). Likewise, *SOCS3* increases in transcript abundance with IFN-γ when compared to control cells (*F*_(2,12)_ = 18.971, p < 0.01) at 10 ng/ml (Tukey HSD, p = 0.005) and 50 ng/ml (Tukey HSD, p < 0.01), but remains unchanged between the two dosages (Tukey HSD, p = 0.124). *IL-10* (*F*_(2,11)_ = 1.305, p = 0.310), *CD200R1* (*F*_(2,12)_ = 1.534, p = 0.255), and *ARG1* (*F*_(2,12)_ = 0.979, p = 0.404) remain unchanged in response to IFN-γ treatment (**Figure 8a-e**).

**Fig. 8.**
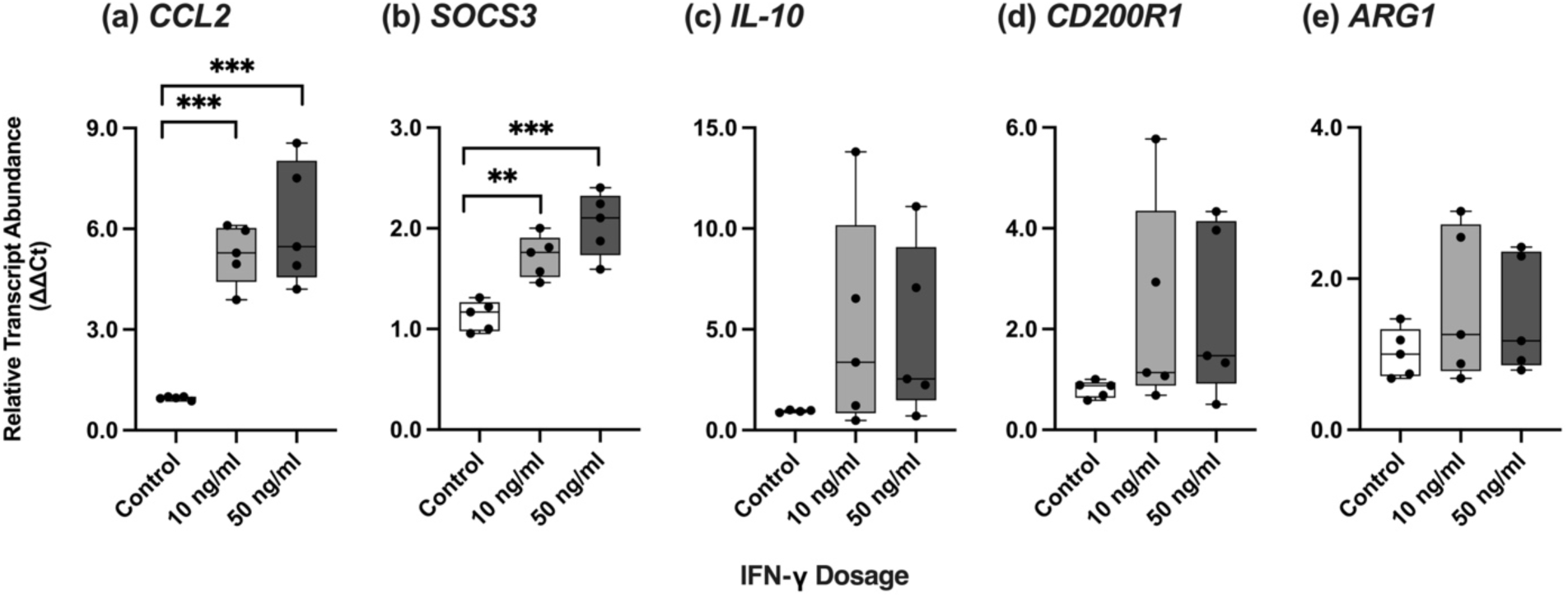
Relative transcript abundance of anti-inflammatory markers in response to IFN-γ (10 ng/ml and 50 ng/ml) in HMC3 cells. **(a)** *CCL2*, **(b)** *SOCS3*, **(c)** *IL-10*, **(d)** *CD200R1*, and **(e)** *ARG1*. Data are mean 2^−ΔΔC*t*^ with individual points (n=5 biological replicates per experimental group). Targets are normalized to the geometric mean of reference genes: *GAPDH, PKM* and *18S*. Significant differences between treatment groups were determined using a one-way ANOVA with Tukey HSD (* p<0.05; ** p<0.01; *** p<0.001).

## Discussion

Microglia play critical roles in the development and maintenance of the CNS from gestation to adulthood (Butovsky & Weiner, 2018; Ransohoff & El Khoury, 2016). Generally, microglia oversee adaptive and innate immune responses triggered by stimuli, stress, and/or disease and produce a spectrum of neuroprotective and/or neurotoxic effects (Gao et al., 2023). There is little consensus on how to define microglia polarization and classify the resulting phenotypes (J. Wang et al., 2023). In the past, microglia were characterized using peripheral macrophage terminology: “M1” (“classic”) and “M2” (“alternative”) (Hu et al., 2014) where “M1” microglia were thought to release pro-inflammatory factors and, if chronically induced, could lead to disease pathologies, while “M2” microglia exert neuroprotective effects by promoting neuronal repair, waste removal and regeneration (Gao et al., 2023; J. Wang et al., 2023). However, this dichotomous way of classifying microglia oversimplifies their plasticity and hinders our understanding of the functional significance and biological roles of microglia in health and disease (Gao et al., 2023; S. Guo et al., 2022; Ransohoff, 2016). Recent advances in single cell transcriptomics in neurodegenerative disease models have described a high degree of spatial and temporal heterogeneity in genes and biomarkers that correspond to polarization states (Friedman et al., 2018; Keren-Shaul et al., 2017; Mathys et al., 2017). The purpose of our study is to utilize the available information on candidate genes that correspond to microglia homeostasis and polarization and quantify their transcript abundance in HMC3 cells in response to IFN-γ treatment. Further, we present detailed methodology in primer design, validation, and testing that can be utilized to quantify these candidate genes by RT-qPCR. We hope to value-add the existing body of literature on HMC3 biomarkers, and designed and tested reference genes (*GAPDH, PKM, 18S, ACTB, PGK1, TKT1, TPI1*), homeostatic (*CD68, TGF-β, IBA1, BIN1, RGS10*), pro-inflammatory (*IL-6, CXCL10, CCL5, SAA, IL-1 β*), and anti-inflammatory targets (*CCL2, SOCS3, IL-10, CD200R1, ARG1*) in control cells and cells primed with 10 and 50 ng/ml of IFN-γ. Out of the seven candidate genes tested as internal controls, we identified three suitable genes, *GAPDH, PKM*, and *18S.* We also find that a spectrum of pro and anti-inflammatory cytokines changes in transcript abundance in response to IFN-γ and no clear demarcation of the “M1” and “M2” phenotypes.

Past studies of microglial morphology have often used both descriptive and stain-based techniques to gather information on the cell body size, ramification degree, and branching pattern of baseline and polarized microglia (Green & Rowe, 2024; Lawson et al., 1990; Robertson, 1899; Tremblay et al., 2015; Walker et al., 2014). The healthy adult CNS contains longitudinally and radially branched and ramified microglia with a large cell body (Hanisch & Kettenmann, 2007). Microglial phenotypic diversity and transformation is associated with the local environment, where smaller, more compact ameboid cells with short/unbranched processes are associated with an injury, insult or stimulus (Morrison et al., 2017; Walker et al., 2014). We found minimal differences in microglial morphology in response to IFN-γ priming across dosage (10 and 50 ng/ml) (**Figure 2a**). Baseline homeostatic cells display radially branched networks with large cell bodies, similar to cells that received 10 or 50 ng/ml IFN-γ. However, as qualitative observations on microglia morphology are not sufficient to define their functional states, we quantified the cell viability and percent live and dead counts. The effects of IFN-γ on HMC3 cell viability were evaluated using an MTT assay.

The redox potential in viable mammalian cells can be measured by quantifying the conversion of MTT reagent to insoluble formazan (Riss et al., 2013). The amount of available formazan is used as a proxy to calculate metabolic activity and viability. The IFN-γ treated HMC3 cells at both 10 and 50 ng/ml show no significant differences compared to untreated controls (**Figure 2b**), suggesting that at the tested concentrations, IFN-γ does not change cellular metabolic activity. A study by Schultzberg et al. (2010) reported similar findings in the HMC3 cell line, where incubation of microglia with 50 ng/ml IFN-γ for 24 hours does not change MTT-determined cell viability, although longer incubations (48 hours and 72 hours) significantly decrease cell viability. Lindberg et al. (2005) also reported no change in cell viability of HMC3 cells incubated with 1-100 ng/ml IFN-γ. While MTT assays offer a straightforward method to assess cell viability, it is important to note their limitations. Factors such as cell seeding density, type of culture media, incubation time, and cytotoxicity of cell treatment conditions may impact absorbance outputs and interpretations (Ghasemi et al., 2021). To minimize potential biases in the interpretation of our results, we supplemented MTT assay findings with live and dead cell counts. We found that IFN-γ treatment, at both dosages, does not change the percentage of live and dead cell counts compared to the untreated controls (**Figure 2c**). As such, we conclude that neither 10 nor 50 ng/ml IFN-γ affects HMC3 cell viability. During polarization, microglia respond to inflammatory challenges by changing their morphology, ultrastructure, motility and function, and gene expression. Microglial molecular profiles are highly heterogeneous and differ across brain regions and developmental stages (Paolicelli et al., 2022). Therefore, it is imperative to select suitable and stable internal controls in gene expression studies to prevent erroneous interpretations (Kozera & Rapacz, 2013; Vandesompele et al., 2002).

We validated seven candidate reference genes involved in glycolysis, molecular transport, and protein translation. Of these, *GAPDH, PKM*, and *18S* (**Figure 3a**) remain unchanged in response to IFN-γ treatment. Analysis of thermal stability also indicates that the genes, from most to least stable, are *GAPDH* > *PKM* > *18S*. *GAPDH* is one of the most cited reference genes in RT-qPCR studies (Barber et al., 2005). GAPDH stability can be attributed to its central, rate determining role in glycolysis (Nicholls et al., 2012; Tristan et al., 2011). PKM also plays a central role in glycolysis, catalyzing the dephosphorylation of phosphoenolpyruvate to pyruvate (C. Liu et al., 2022). *PKM* is also a widely used reference gene (C. Liu et al., 2022). Our findings on the suitability of *GAPDH* and *PKM* as internal references in HMC3 cells is corroborated by Hernández-Ochoa et al. (2021) who tested ten reference genes in the HMC3 cell line. They found that *PKM* is the most stable gene, with a thermal stability of 0.009 and *GAPDH*, while not the most stable among the candidates, was still suitable, with a thermal stability of 0.020. A study by Fazzina et al. (2024) in HMC3 cells post 24-hour IFN-γ (1 μg/ml) and glucose (5 g/l) treatments, identified *GAPDH* to be a suitable reference gene, with a thermal stability of 0.315. *18S*, an RNA component of the 40S ribosomal subunit, is another popular reference gene used for normalization due to its ubiquitous expression across species and cells (Barta et al., 2023; Kuchipudi et al., 2012). Our findings suggest that while *18S* transcript levels do not change with IFN-γ treatment, it is not as thermally stable as *GAPDH* and *PKM*. The same study by Fazzina et al. (2024) found similar results for *18S* in HMC3 cells, with a thermal stability of 0.230. Transcript levels of *ACTB, PGK1, TKT1*, and *TPI1* significantly change in response to IFN-γ (**Figure 3b**).

ACTB is an essential component of the cytoskeleton and mediates cell motility and division (Bunnell et al., 2011; Nimmerjahn et al., 2005). Given its integral role in cell structure and migration, *ACTΒ* is highly expressed in microglia, and thus often used as a reference gene (Gosselin et al., 2017). Despite this, we found that *ACTΒ* transcript levels change with 10 and 50 ng/ml IFN-γ. On the contrary, a study by Straccia et al. (2011) using murine primary microglia reported no change in transcript abundance of *ActΒ* after a 6-hour incubation with 100 ng/ml LPS combined with 0.1 ng/ml IFN-γ. Fazzina et al. (2024) found that 24 hour incubation of HMC3 cells with 1 μg/ml IFN-γ and 5 g/l glucose also did not change transcript abundance of *ACTB*, although the reported thermal stability was 0.378. The differences across studies can be attributed to the type and dosage of the pro-inflammatory stimulus used for priming HMC3 cells. For example, in the study by Straccia et al. (2011) primary microglia were harvested and cultured from mouse cerebral cortices at postnatal day 1, compared to the HMC3 cells used in our study which are immortalized human embryonic microglia. Furthermore, the dosage and type of stimulus was different.

TPI1 and PGK1 are involved in the glycolytic/gluconeogenic pathways, catalyzing the interconversion of dihydroxyacetone phosphate and glyceraldehyde-3-phosphate, and catalyzing the formation of 3-phosphoglycerate and ATP from 1,3-diphosphoglycerate, respectively (Jiang et al., 2017; Li et al., 2016). Hernández-Ochoa et al. (2021) found that *TPI1* did not change in untreated HMC3 cells or pediatric brain glioma biopsies, with a thermal stability of 0.009. This is perhaps unsurprising, given they did not use an acute polarization stimulus. Further, metabolic profiles of tumor cells cannot easily be compared to non-tumor, immortalized cells. Similar to our results, which indicate an increase in *PGK1* transcript abundance with IFN-γ treatment, Romero-Ramírez et al. (2022) reported an increase in *Pgk1* transcript abundance with LPS (10 ng/ml, 24 hours) treatment compared to controls in murine microglia.

TKT1 is involved in the pentose phosphate pathway and facilitates the glycolytic conversion of pentose sugars into triose and hexose sugars and to generate ribose-5-phosphate *de novo* for nucleic acid synthesis (Ricciardelli et al., 2015). The same study by Hernández-Ochoa et al. (2021) found that in untreated HMC3 cells and pediatric brain glioma, *TKT1* transcript abundance remained unchanged, with a thermal stability of 1.689. These findings emphasize that the outcome of gene expression studies in HMC3 cells vary significantly depending on the stimulus, dosage and the nature of the cells. We cannot assume a reference gene that was used in one study will be suitable for a second study with different parameters. As such, it is imperative that a combination of stable reference candidates is used for normalization, versus using a single, internal control (Vandesompele et al., 2002).

Under homeostatic conditions, microglia maintain CNS function, including immunological surveillance, synaptic pruning and the production of neurotrophic factors (Lima et al., 2022). Specifically, microglia use rounded motile protrusions extending out of their cell body to sense changes in their microenvironment (Nimmerjahn et al., 2005). Of the five homeostatic markers we tested in our study, *CD68* increases and *TGF-β* decreases in transcript abundance with IFN-γ treatment (**Figure 5a-b**), while *IBA1, BIN1*, and *RGS10* remain unchanged (**Figure 5b-d**).

CD68 is listed as a homeostatic marker in our study because it is a transmembrane glycoprotein associated with the endo-lysosomal compartment and mediates antigen processing and peptide transport (Chistiakov et al., 2017). CD68 also controls microglial responses to tissue damage. This means that CD68 is typically upregulated during pro-inflammation at the gene and protein levels (Hendrickx et al., 2017; Jurga et al., 2020). Similar to our findings, Wong et al. (2005) reported that after 24 hour incubation with 10 ng/ml IFN-γ and 100 ng/ml LPS concurrently, *Cd68* gene expression was upregulated in murine BV-2 microglia. On the contrary to the increase in *CD68* levels, we saw a decrease in transcription abundance of TGF-β.

TGF-β promotes microglial proliferation (Bureta et al., 2020), controls the homeostasis of immune cells (Sheng et al., 2015; Spittau et al., 2020) and reduces chemokine, cytokine, and reactive oxygen species (ROS) production in microglia (Rustenhoven et al., 2016). A study by Mitchell et al. (2014) found that LPS treatment to primary rat microglia downregulates TGF-β signaling and suppresses its ability to induce cell death and express pro-inflammatory cytokines. They proposed that other pro-inflammatory factors can inhibit TGF-β signaling to maintain a pro-inflammatory phenotype. As such, the IFN-γ induced decrease in *TGF-β* levels combined with no changes in cell viability or live and dead cell counts, could be a biological outcome of pro-inflammation. However, protein analysis is required to delineate the exact mechanisms.

IBA-1 is a calcium-binding protein that has actin-bundling capability and participates in membrane ruffling and phagocytosis (Ohsawa et al., 2004). IBA-1 is found in infiltrating macrophages and is expressed in neuroglia, and absent in neurons, making it a popular biomarker for neuroglia studies (Hopperton et al., 2018; Jurga et al., 2020). However, Ito et al. (1998) found that IBA-1 identifies all microglia phenotypes, regardless of polarization state. Thus, like our findings, they found that *IBA-1* transcript abundance typically does not change with polarization. A study by Hendrickx et al. (2017) also reported that *IBA-1* is a suitable biomarker for structural studies in the absence of pathology, and that it is not always concurrently expressed with other homeostatic markers such as *CD68.* Furthermore, under most conditions, the total number of microglia in the CNS does not dramatically change (Ginhoux et al., 2013; Lawson et al., 1992) and a reduction in the total number of microglia may cause the existing population to differentiate to replace the lost cells (Butovsky and Weiner, 2018).

BIN1 is involved in membrane remodeling, cytoskeletal maintenance, and regulation of the cell cycle (Sudwarts et al., 2022). Similar to our findings of stable expression in response to IFN-γ, a study by Sudwarts et al. (2022) found that LPS treatment did not change *Bin1* expression or localization in murine primary neonatal microglia. Interestingly, they found that BIN1 positively regulates the transcription of *Sfpi1* and *Irf1* regulators, which in turn control numerous microglial genes under both homeostatic and pro-inflammatory conditions. This could potentially explain why *BIN1* levels remain unchanged with pro-inflammatory stimulus.

RGS10 belongs to a family of GTPase accelerating proteins and is involved in negatively regulating G-protein coupled receptor signaling by increasing GTP hydrolysis (Lee et al., 2013). Wendimu et al. (2021) found that *Rgs10* knockout cells had dysregulated pro-inflammatory and anti-inflammatory responses. Furthermore, they found that 10 ng/ml of LPS treatment (for 24 hours) to murine BV-2 microglia caused *Rgs10* to downregulate the expression of select pro-inflammatory mediators, including cyclooxygenase 2 and its primary metabolic product, prostaglandin E2. Taken together, these studies may suggest that minimal changes in *RGS10* in response to IFN-γ we reported, may be due to its cytoprotective role in the regulation of microglial functional states rather than only a pro-inflammatory state. Therefore, *RGS10* could exhibit a higher level of baseline transcript abundance in HMC3 that warrants a further increase in response to stimuli and/or stress.

The disruption to the homeostatic CNS environment triggers a morphological shift in microglia and leads to the increased production of intracellular molecules and surface antigens (Perry and Holmes, 2014). During inflammation, microglia can activate and recruit bone marrow-derived macrophages and monocytes to the site of injury (Pallarés-Moratalla and Bergers, 2024). Additionally, polarized microglia produce more IL-1, TNF-α, IL-6, and nitric oxide (Combrinck et al., 2002; Cunningham et al., 2009, 2005; Perry and Holmes, 2014). Correspondingly, our study found that the pro-inflammatory target *IL-6* increases in transcript abundance with IFN-γ (**Figure 7a**). However, the increased levels of *IL-6* are not a clear indication that the HMC3 cells have polarized into a pro-inflammatory state, because IL-6 exhibits anti or pro-inflammatory properties depending on the activation ligand and upstream receptor complex (Tilg et al., 1997). Thus, IL-6 can have neuroprotective effects, but elevated IL-6 levels in inflammation and injury can also decrease neuron survival (Kummer et al., 2021).

Microglia can recruit peripheral immune cells by secreting chemokines such as CXCL10 and CCL5 (Jurga et al., 2020; Kim and Joh, 2006). Our study found that compared to the baseline cells, the transcript abundance of *CXCL10* and *CCL5* robustly increases with IFN-γ treatment (**Figure 7b-c**). Previous work has also found that IFN-γ stimulates increased production of *Cxcl10* in murine microglia (Vanguri and Farber, 1994) and pro-inflammatory stimuli are known to increase CCL5 production (Pittaluga, 2017). The *CXCL10* receptor, CXCR3, is expressed in primary human microglia (Biber et al., 2002), and CXCL10 is a gene that is highly expressed in neurodegenerative diseases (Rappert et al., 2004). Therefore, the lack of *CXCL10* mRNA in baseline HMC3 cells and increased abundance in response to IFN-γ treatment demonstrates the importance of CXCL10 upregulation during polarization.

The remaining pro-inflammatory markers, *SAA* and *IL-1β*, used in our study are biologically connected. A study by Yu et al. (2019) in a murine model showed that SAA can activate the NLRP3 inflammasome and increase IL-1β production. SAA also induces a pro-inflammatory phenotype in microglia (Yu et al., 2019). We find that *SAA* transcript abundance increases with 10 ng/ml of IFN-γ treatment (**Figure 7d**), which aligns with previous findings that the production of SAA, an acute phase protein, is triggered by pro-inflammatory cytokines, like IFN-γ (Yu et al., 2019). Further, we found that *IL-1β* expression is not sensitive to 10 or 50 ng/ml of IFN-γ treatment (**Figure 7e**). The lack of changes may indicate that, unlike LPS, which universally increases *IL-1β* levels in microglia (Cacci et al., 2008; He et al., 2021; Ishijima et al., 2021; Olajide et al., 2013), IFN-γ may produce individual responses that are species, stress, and dosage specific. For instance, IFN-γ (1, 10, and 100 ng/ml) directly inhibits *IL-1β* transcription in murine bone marrow–derived macrophages and dendritic cells (Eigenbrod et al., 2013).

During inflammation, polarized microglia can also produce anti-inflammatory cytokines, growth factors, and neurotrophic factors to stimulate phagocytosis and support tissue repair and neuroglia survival (Guo et al., 2022). The chemokine CCL2 can induce anti-inflammatory polarization of macrophages (Roca et al., 2009) and is reported to be secreted mostly by “M2” microglia (Jurga et al., 2020). Therefore, we have classified CCL2 as an anti-inflammatory marker, although we recognize that CCL2 can exhibit pro-inflammatory properties (Laffer et al., 2019). Our study finds that *CCL2* transcript abundance robustly increases with IFN-γ treatment (**Figure 8a**), which is likely due to its role in recruiting peripheral immune cells to the site of injury (Cherry et al., 2020; Sozzani et al., 1994). The receptor of CCL2, CCR2, is expressed in rat microglia, but the CCL2/CCR2 axis does not uniquely contribute to inflammatory modulation (Conductier et al., 2010). A study by Pedragosa et al. (2020) found that CCR2^+^ monocytes contributed to the expression of genes involved in angiogenesis and repair in a murine model. As such, we can only conclude that CCL2 upregulation is likely due to microglia polarization but cannot comment on the anti or pro-inflammatory direction of the transformation.

SOCS3 inhibits STAT activation via JAK-STATs, specifically the regulation of STAT3 activation in response to cytokines using the gp130 receptor, like IL-6 (Carow and Rottenberg, 2014; Van Wagoner and Benveniste, 1999). Since increased SOCS3 is important in reducing pro-inflammation (Chakrabarti et al., 2018), we have categorized it as an anti-inflammatory marker of microglia in this study. We found an increase in *SOCS3* transcript abundance with IFN-γ treatment (**Figure 8b**), which could be a compensatory, anti-inflammatory response to the increased *IL-6* levels seen in our study (**Figure 7a**).

However, the anti-inflammatory markers *IL-10*, *CD200R1* and *ARG1* did not change in transcript abundance with IFN-γ (**Figure 8c-e**). IL-10, an anti-inflammatory cytokine, can modulate glial activation and regulate cytokine levels in the CNS (Laffer et al., 2019). Increased IL-10 secretion is generally associated with anti-inflammatory outcomes (Henkel et al., 2009). The CD200R1/CD200 axis in microglia is involved in inhibiting the expression of pro-inflammatory molecules like IL-6 (Hernangomez et al., 2014; Rabaneda-Lombarte et al., 2021), and inhibits pathways involved in microglial activation, including p38, extracellular signal-regulated kinase (ERK), and c-Jun terminal kinase (JNK) (Manich et al., 2019; Zhao et al., 2019). In addition, CD200R1 inhibits pro-inflammatory polarization (Rabaneda-Lombarte et al., 2020). ARG1 hydrolyzes arginine to ornithine and urea (Wu and Morris, 1998), thereby competing with nitric oxide synthase (NOS) and decreasing nitric oxide (NO) production as a result (Caldwell et al., 2015). *ARG1* is often classified as an “M2” marker of microglia and macrophages (Hu et al., 2012). Given the pro survival roles these proteins play in neuroglia, the lack of transcriptional changes in response to 10 and 50 ng/ml of IFN-γ is unexpected. The lack of differential expression is likely attributed to technical difficulties in measuring these targets across baseline and polarized HMC3 cells. Specifically, we found that *IL-10*, *CD200R1* and *ARG1* expression is tightly distributed in the control cells but varies largely in cells treated with 10 and 50 ng/ml of IFN-γ. Interestingly, select biological replicates were more responsive to IFN-γ compared to others. For example, one biological replicate illustrated the lowest *IL-10, CD200R1* and *ARG1* abundance in the 50 ng/ml IFN-γ-treated cells. The same was observed for the biological replicate that illustrated the highest transcript abundance for the three targets, however they were not statistical outliers. Further, this distribution is unique to *IL-10, CD200R1* and *ARG1* and is absent in other targets. Our results highlight the complex heterogeneity of the HMC3 polarization spectrum to stimuli and emphasizes the importance of using a comprehensive set of biomarkers to characterize HMC3 morphology and function. HMC3 cells likely do not exhibit polarization states. Rather, it is a dynamic polarization spectrum. Moreover, the gene expression patterns of biomarkers largely depend on the stimulus, duration, dosage, cell line and developmental stage.

## Conclusion

The immortalized HMC3 cell line enables the study of microglia dynamics and the polarization spectrum in response to a variety of stimuli and stress and is an ideal platform for pharmacological screening. It is a powerful tool that complements *in vivo* microglia research and retains cellular and morphological characteristics to those of primary microglia. In our study, we have provided detailed methodology on biomarker selection, primer design, validation parameters, and quantification of homeostatic, anti-inflammatory, and pro-inflammatory targets, in response to IFN-γ treatment. Further, our study analyzed the transcript abundance of a set of biomarkers that can be used to characterize HMC3 polarization as a spectrum, instead of states, and improves our understanding of how microglia respond to a widely used cytokine, IFN-γ. Our findings also illustrate that microglial responses to stress are often unique and depend on the type of stimulus (*e.g.* dosage, duration) and the cell-type (*e.g.* culture age, organism, developmental stage). Ultimately, our findings contribute to a greater understanding of microglia polarization and increase the rigor of gene expression analysis and biomarker selection in the HMC3 cell line.

## Data Availability Statement

The authors confirm that the data supporting the findings of this study are available within the article and its supplementary materials.

## Acknowledgements

We like to thank Dr. Patrick O. McGowan (Ph.D.) from the University of Toronto for providing a vial of the HMC3 cell line for propagation and use in the Wijenayake Lab at The University of Winnipeg. We also like to thank the support staff at The University of Winnipeg for their assistance.

