## Supplementary Figure S1 for "The glia club: Validation of polarization biomarkers for human microglia (HMC3) using quantitative real time RT-qPCR"

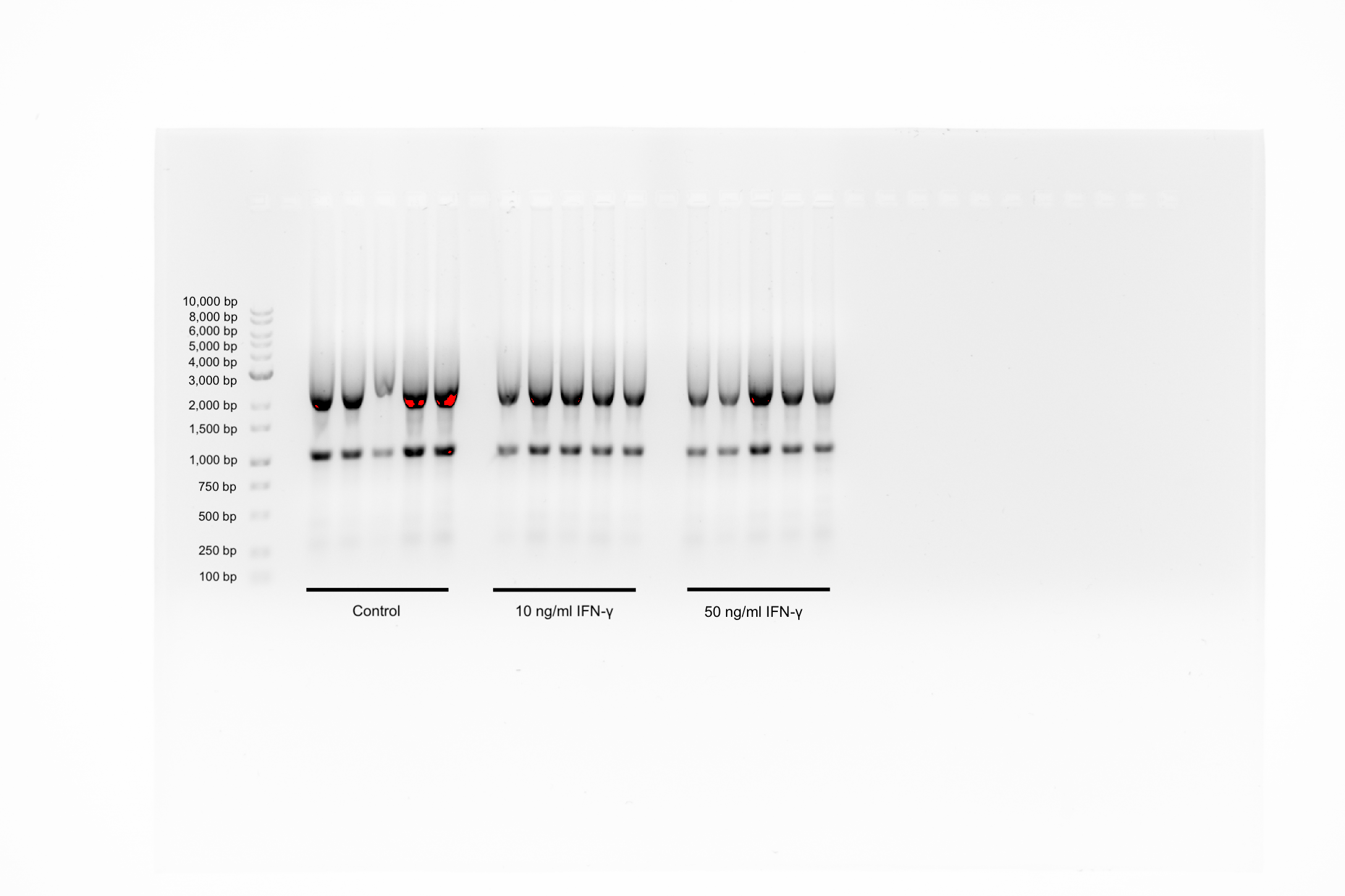

**Supplementary Figure S1.** 1% TAE-agarose gel to verify RNA stability and integrity. The ladder range is from 100 bp – 10,000 bp. RNA profiles for each group (control, 10 ng/ml IFN-γ, and 50 ng/ml IFN-γ) show distinct 5S, 18S, and 28S subunit distribution.
