## Supplementary Figure S2 for "The glia club: Validation of polarization biomarkers for human microglia (HMC3) using quantitative real time RT-qPCR"

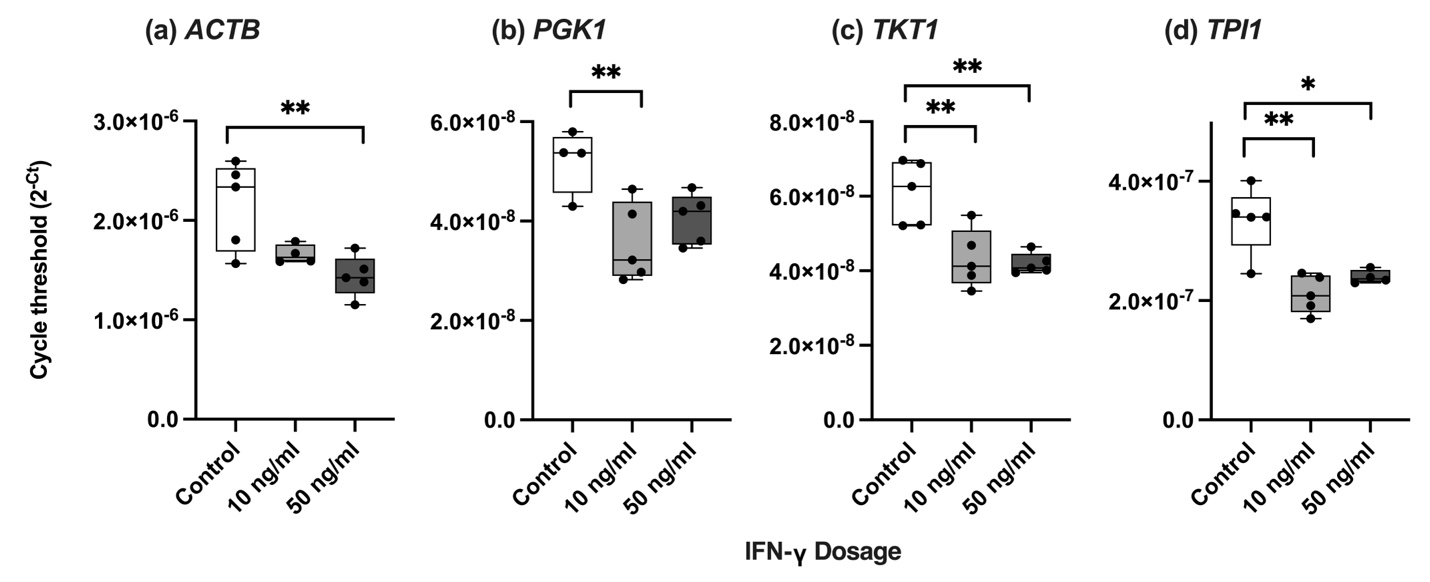

**Supplementary Figure S2** Relative transcript abundance of four candidate reference genes with differential expression in response to interferon-gamma (IFN-γ) treatment (10 ng/ml and 50 ng/ml) when compared to untreated control cells. **(a)** *ACTB*, **(b)** *PGK1*, **(c)** *TKT1*, and **(d)** *TPI1*. Data are mean $2^{-Ct}$with individual points (n=5 biological replicates per experimental group). 2 x 10^5^ cells were plated per well. Significant differences between treatment groups were determined using a one-way ANOVA with Tukey HSD (* p < 0.05; ** p < 0.01; *** p < 0.001).
