## Supplementary Table S1 for "The glia club: Validation of polarization biomarkers for human microglia (HMC3) using quantitative real time RT-qPCR"

**Supplementary Table S1.** Primer sequences and RT-qPCR parameters for targets that were not further tested. Targets are identified by gene name, NCBI accession number, targeted transcript variants, forward and reverse primer sequences, annealing temperature (°C), cDNA amount (ng), threshold cycle (Ct), product size (bp), and function. The sources of the primers (designed using NCBI or obtained from literature) are indicated. All primers target *Homo sapiens*.

| **Gene** | **NCBI Accession #** | **Transcript variants** | **Forward primer (5’ – 3’)** | **Reverse primer (5’ – 3’)** | **Annealing temperature (℃)** | **cDNA loaded (ng)** | **Product size (bp)** | **Function** | **Source of primer** |
| --- | --- | --- | --- | --- | --- | --- | --- | --- | --- |
| *RGS10* | NM_002925.4 | 1 & 2 | CGGCATCCCTGGAGAATCTG | ATCAGAGGGTGCGGTTCTTC | 60 | 10 | 262 | RGS proteins function as GTPase-activating proteins (GAPs). Regulate Gi- and Gq-mediated signaling from many mammalian GPCRs (Larminie et al., 2004). | Designed |
| *CXCL10* | NM_001565.4 | 1 | GTGGCATTCAAGGAGTACCTC | TGATGGCCTTCGATTCTGGATT | 62 | 10 | 198 | Chemokine. CXCL10/CXCR3-A axis involved in chemotaxis and proliferation of cell types, CXCL10/CXCR3-B induces apoptosis (Bodnar et al., 2006; Kelsen et al., 2004; Lasagni et al., 2003). | (Peferoen et al., 2015) |
| *CCL5* | NM_002985.3 | 1 & 2 | GAGTATTTCTACACCAGTGGCAAG | TCCCGAACCCATTTCTTCTCT | 61 | 10 | 104 | Pro-inflammatory chemokine: chemotaxis and immune response (Yu-Ju Wu et al., 2020). | (Yu-Ju Wu et al., 2020) |
| *CCL2* | NM_002982.4 | N/A | TGCAATCAATGCCCCAGTCA | GGGTCAGCACAGATCTCCTT | 60 | 10 | 156 | Binds to CCR2; can recruit and activate monocytes, neutrophils, and lymphocytes (Bose & Cho, 2013). | (Zhang et al., 2022) |
