## Supplementary Table S2 for "The glia club: Validation of polarization biomarkers for human microglia (HMC3) using quantitative real time RT-qPCR"

**Supplementary Table S2.** Bioinformatics of working primers that were not further tested. Targets are identified by gene name, NCBI accession number, targeted transcript variants, and forward and reverse primer sequences. All primers target *Homo sapiens.* Product size (bp), forward and reverse melting temperatures (°C) and GC content (%), hairpin temperature (°C), and homodimer and heterodimer values (%) are listed. The sources of the primers (designed using NCBI or obtained from literature) are indicated.

| **Gene** | **NCBI Accession #** | **Forward primer (5’ – 3’)** | **Reverse primer (5’ – 3’)** | **Product size (bp)** | **Melting temperature forward (℃)** | **Melting temperature reverse (℃)** | **GC content forward (%)** | **GC content reverse (%)** | **Hairpin temperature (℃)** | **Homodimer (%)** | **Heterodimer (%)** | **Source of primer** |
| --- | --- | --- | --- | --- | --- | --- | --- | --- | --- | --- | --- | --- |
| *RGS10* | NM_002925.4 | CGGCATCCCTGGAGAATCTG | ATCAGAGGGTGCGGTTCTTC | 262 | 64.1 | 63.8 | 60.0 | 55.0 | 28.3 | 11.4 | 18.9 | Designed |
| *CXCL10* | NM_001565.4 | GTGGCATTCAAGGAGTACCTC | TGATGGCCTTCGATTCTGGATT | 198 | 62.3 | 64.1 | 52.4 | 45.5 | 48.0 | 12.3 | 15.3 | (Peferoen et al., 2015) |
| *CCL5* | NM_002985.3 | GAGTATTTCTACACCAGTGGCAAG | TCCCGAACCCATTTCTTCTCT | 104 | 62.9 | 63.2 | 45.8 | 47.6 | 37.2 | 12.2 | 12.2 | (Yu-Ju Wu et al., 2020) |
| *CCL2* | NM_002982.4 | TGCAATCAATGCCCCAGTCA | GGGTCAGCACAGATCTCCTT | 156 | 64.7 | 63.4 | 50.0 | 55.0 | 54.0 | 17.6 | 15.3 | (Zhang et al., 2022) |
