## Supplementary Table S3 for "The glia club: Validation of polarization biomarkers for human microglia (HMC3) using quantitative real time RT-qPCR"

**Supplementary Table S3.** Bioinformatics of non-working primers. Targets are identified by gene name, NCBI accession number, targeted transcript variants, and forward and reverse primer sequences. All primers target *Homo sapiens.* Product size (bp), forward and reverse melting temperatures (°C) and GC content (%), hairpin temperature (°C), and homodimer and heterodimer values (%) are listed. The sources of the primers (designed using NCBI or obtained from literature) are indicated.

| **Gene** | **NCBI Accession #** | **Forward primer (5’ – 3’)** | **Reverse primer (5’ – 3’)** | **Product size (bp)** | | **Melting temperature forward (℃)** | **Melting temperature reverse (℃)** | **GC content forward (%)** | **GC content reverse (%)** | **Hairpin temperature (℃)** | **Homodimer (%)** | **Heterodimer (%)** | **Source of primer** |
| --- | --- | --- | --- | --- | --- | --- | --- | --- | --- | --- | --- | --- | --- |
| *ARG1* | NM_001244438.2 | TCTTGGAGTCATCTGGGTGGA | TCCTGGCACATCGGGAATCT | 133 | 64.6 | | 65.3 | 52.4 | 55.0 | 26.4 | 5.0 | 11.4 | Designed |
| *BIN1* | NM_139343.3 | CCTGCTGTGGATGGATTACC | GCTTTCTCAAGCAGCGAGAC | 219 | 62.2 | | 63.2 | 55.0 | 55.0 | 29.1 | 8.2 | 21.6 | (De Rossi et al., 2016) |
| *CCL17* | NM_002987.3 | CTCCAGGGATGCCATCGTTT | CCTCTCAAGGCTTTGCAGGTA | 109 | 64.3 | | 64.0 | 55.0 | 52.4 | 42.6 | 12.2 | 15.2 | Designed |
| *CCL18* | NM_002988.4 | CTGCCCAGCATCATGAAGG | CCTCAGGCATTCAGCTTCAG | 186 | 62.8 | | 62.6 | 57.9 | 55.0 | 34.5 | 22.4 | 21.3 | (Yang et al., 2023) |
| *CCL18* | NM_002988.4 | CTCTGCTGCCTCGTCTATACC | TTAGGAGGATGACACCTGGCT | 103 | 63.3 | | 64.6 | 57.1 | 52.4 | 16.6 | 9.3 | 15.8 | Designed |
| *CCL22* | NM_002990.5 | CGTGATTACGTCCGTTACCG | AAGGTTAGCAACACCACGC | 98 | 62.3 | | 62.9 | 55.0 | 52.6 | 37.6 | 15.9 | 17.4 | (Peferoen et al., 2015) |
| *CCL22* | NM_002990.5 | ATGGATCGCCTACAGACTGC | GGATCGGCACAGATCTCCTT | 236 | 63.6 | | 63.4 | 55.0 | 55.0 | 36.1 | 12.2 | 16.0 | Designed |
| *CCL5* | NM_002985.3 | ACCAGTGGCAAGTGCTCCA | ACCCATTTCTTCTCTGGGTTGGCA | 86 (variant 1)  168 (variant 2) | 66.2 | | 68.1 | 57.9 | 50.0 | 43.2 | 13.5 | 13.9 | (Liu et al., 2014) |
| *CCR2* | NM_001123041.3 | CGAGAGCGGTGAAGAAGTCA | AGCATGTTGCCCACAAAACC | 144 | 63.6 | | 64.1 | 55.0 | 50.0 | 31.2 | 9.5 | 11.0 | Designed |
| *CD11B* | NM_000632.4 | CGCAATGACCTTCCAAGAGAAC | CTGTAGTCGCACTGGTAGAGG | 138 | 63.4 | | 63.1 | 50.0 | 57.1 | 32.0 | 8.6 | 12.0 | Designed |
| *CD11B* | NM_001145808 | CACGGATGGAGAAAAGTTTGG | TGGATGCGATGGTATTAAGCTC | 149 | 61.4 | | 62.3 | 47.6 | 45.5 | 47.0 | 18.8 | 11.7 | (Roberts & Robinson, 2014) |
| *CD163* | NM_001370145.1 | GAAGACAGAGACAGCGGCTT | GTGTGGCTCAGAATGGCCT | 140 | 65.0 | | 65.4 | 55.0 | 57.9 | 16.0 | 12.6 | 16.5 | Designed |
| *CD163* | NM_001370145.1 | GAGTCCCTTCACCATTACTGTG | GACTTTCACTTCCACTCTCCC | 133 | 63.2 | | 63.9 | 50.0 | 52.4 | 30.6 | 8.6 | 11.8 | (Asahchop et al., 2017) |
| *CD163* | NM_001370145.1 | TTTGTCAACTTGAGTCCCTTCAC | TCCCGCTACACTTGTTTTCAC | 127 | 63.0 | | 62.8 | 43.5 | 47.6 | 22.1 | 13.6 | 9.7 | (Sneeboer et al., 2019) |
| *CD200R1* | NM_138806.4 | ATGCTCTGCCCTTGGAGAAC | TTGTTTGGTTGAGCAGCACC | 95 | 65.3 | | 64.8 | 55.0 | 50.0 | 42.2 | 8.1 | 21.1 | Designed |
| *CD200R1* | NM_138806.4 | ATCTTCTTAGTGGCCGAAGC | GCACAGCATTTGTAGCCATC | 193 | 63.0 | | 62.9 | 50.0 | 50.0 | 38.1 | 24.0 | 21.1 | (Rabaneda-Lombarte et al., 2022) |
| *CD206* | NM_002438.4 | CGTTCCTTTGGACGGATGGA | CCTCGTTTACTGTCGCAGGT | 172 | 65.1 | | 64.9 | 55.0 | 55.0 | 37.1 | 12.0 | 12.0 | Designed |
| *CD206* | NM_002438.4 | GACTCCCGAACCCAAATGTCC | TCGCCATATTGTTTGCTGTTCC | 184 | 65.6 | | 64.8 | 57.1 | 45.5 | 24.7 | 8.6 | 13.6 | (Zeiner et al., 2019) |
| *CD68* | NM_001040059 | CATCTCTGTACTGAACCCCAAC | CCATGTAGCTCAGGTAGACAAC | 149 | 62.2 | | 62.0 | 50.0 | 50.0 | 22.7 | 9.1 | 29.9 | (Asahchop et al., 2017) |
| *CRP* | NM_001329057.2 | CAAAGCCTTCACTGTGTGCC | CCAGAACTCCACGATCCCTG | 241 | 60.0 | | 59.8 | 55.0 | 60.0 | 32.6 | 9.2 | 9.1 | Designed |
| *CRP* | NM_001329057.2 | TTTTCTCGTATGCCACCAAG | TTTCCAATGTCTCCCACCAG | 322 | 56.1 | | 57.4 | 45.0 | 50.0 | 18.1 | 9.5 | 5.1 | (Gould & Weiser, 2001) |
| *CXCL13* | NM_006419.3 | GCTTGAGGTGTAGATGTGTCC | CCCACGGGGCAAGATTTGAA | 83 | 62.2 | | 65.0 | 52.4 | 55.0 | 18.5 | 8.7 | 12.7 | (Jiang et al., 2016) |
| *CXCL13* | NM_006419.3 | CCAAGGTGTTCTGGAGGTCT | TTGAGGGTCCACACACACAAT | 181 | 63.3 | | 64.2 | 55.0 | 47.6 | 33.7 | 13.5 | 17.5 | Designed |
| *IBA1* | D86438.1 | ACCAGGGATTTACAGGGAGGA | CCAGTTTGGAGGGCAGATCC | 133 | 64.5 | | 64.4 | 50.0 | 60.0 | 33.6 | 7.5 | 15.0 | Designed |
| *IL-10* | NM_001382624.1 | CTGAGAACCAAGACCCAGACA | TGTCAAACTCACTCATGGCT | 195 | 64.4 | | 62.8 | 52.0 | 45.0 | 30.1 | 9.4 | 13.6 | Designed |
| *IL-10* | NM_001382624.1 | GATCTCCGAGATGCCTTCAGC | GGAAGAAATCGATGACAGCGC | 226 | 65.1 | | 64.5 | 57.1 | 52.4 | 44.5 | 15.3 | 12.5 | (Curto et al., 2004) |
| *IL-12B* | NM_002187.3 | GTCACAAAGGAGGCGAGGTT | ACTGATTGTCGTCAGCCACC | 179 | 64.3 | | 64.3 | 55.0 | 55.0 | 0.0 | 9.2 | 15.8 | (Xu et al., 2016) |
| *IL-13* | NM_002188.3 | CATGGCGCTTTTGTTGACCA | AGCTGTCAGGTTGATGCTCC | 181 | 64 | | 64.1 | 50.0 | 55.0 | 34.3 | 24.2 | 11.9 | (Pallio et al., 2021) |
| *IL-13* | NM_001354993.2 | AGGATGCTGAGCGGATTCTG | AAACTGGGCCACCTCGATTT | 93 | 63.9 | | 64.4 | 55.0 | 50.0 | 21.0 | 12.0 | 11.3 | Designed |
| *IL-18* | NM_001243211.2 | TGACCAAGGAAATCGGCCTC | GCCATACCTCTAGGCTGGCT | 117 | 64.2 | | 65.2 | 55.0 | 60.0 | 30.9 | 22.6 | 19.0 | Designed |
| *IL-4* | NM_001354990.2 | GAACAGCCTCACAGAGCAGA | CTTTGTAGGCGAGTCGAGGT | 162 | 64.7 | | 64.4 | 55.0 | 55.0 | 34.9 | 8.8 | 20.3 | Designed |
| *IL-4* | NM_001354990.2 | AACAGCCTCACAGAGCAGAAGAC | GTGTTCTTGGAGGCAGCAAAG | 72 | 67 | | 64.7 | 52.0 | 52.4 | 34.7 | 7.7 | 22.9 | (Narantuya et al., 2010) |
| *P2RY12* | NM_022788.5 | TGCCCGAATTCCTTACACCC | TTTGGGTCACCACCATCCTG | 249 | 64.2 | | 64.2 | 55.0 | 55.0 | -2.1 | 20.4 | 17.9 | Designed |
| *P2RY12* | NM_022788.5 | TTTGTGTGTCAAGTTACCTCCG | CTGGTGGTCTTCTGGTAGCG | 101 | 62.7 | | 63.8 | 45.5 | 60.0 | 32.1 | 9.9 | 13.6 | (Murai et al., 2020) |
| *P2RY12* | NM_022788.5 | GTGTCAAGTTACCTCCGTCATA | TAAATGGCCTGGTGGTCTT | 104 | 61.6 | | 61.3 | 45.5 | 47.4 | 14.2 | 9.6 | 11.8 | (Banerjee et al., 2020) |
| *P2RY13* | NM_176894 | TGACTGCCGCCATAAGAA | CTGTGGTGTTCATTGCTTCC | 75 | 61.1 | | 61.4 | 50.0 | 50.0 | 22.7 | 10.1 | 13.7 | (Yi et al., 2014) |
| *P2RY13* | XM_006713664.1 | TGCCGCCATAAGAAGACAGA | GCACCGCTCAGATCTGTTGA | 103 | 63.5 | | 64.3 | 50.0 | 55.0 | 3.0 | 9.2 | 16.6 | Designed |
| *SALL1* | NM_002968.3 | CCTGCGTCTAATCCACTTCTAC | CCAAGGCAGACAAGGAGTTTA | 112 | 61.9 | | 62.1 | 50.0 | 47.6 | 8.0 | 9.2 | 17.1 | (Banerjee et al., 2020) |
| *SALL1* | NM_002968.3 | ATTTGGCGGCAAGATCAGGA | TTTCCCGTCAGCCCACTAAC | 184 | 64.4 | | 64.1 | 50.0 | 55.0 | 30.7 | 11.1 | 16.0 | Designed |
| *TMEM119* | NM_181724.3 | GCCACCCAGAACCTCAAGTC | CTGGGCTTCCTGGGCTAAC | 258 | 64.5 | | 64.2 | 60.0 | 63.2 | 17.1 | 8.1 | 24.4 | Designed |
| *TMEM119* | NM_181724.3 | AGTCCTGTACGCCAAGGAAC | GCAGCAACAGAAGGATGAGG | 92 | 63.7 | | 62.8 | 55.0 | 55.0 | 42.6 | 16.4 | 16.4 | (Washer et al., 2022) |
| *TMEM119* | NM_181724.3 | GTCCACAATATTCGTCAGTC | CTGGTGCATTATATCTCAGC | 149 | 58.1 | | 58.0 | 45.0 | 45.0 | -18.7 | 23.2 | 14.5 | (Lopez-Lengowski et al., 2021) |
| *TMEM119* | NM_181724.3 | CTGCTGATGTTCATCGTCTGT | TCACTCTGGTCCACGTACT | 107 | 62.3 | | 61.9 | 47.6 | 52.6 | 34.0 | 13.7 | 13.6 | (Banerjee et al., 2020) |
| *TMEM119* | NM_181724.3 | CTTCCTGGATGGGATAGTGGAC | GCACAGACGATGAACATCAGC | 96 | 63.4 | | 63.4 | 54.5 | 52.4 | 38.8 | 11.4 | 12.2 | (Han et al., 2018) |
| *TMEM119* | NM_181724.3 | GGACTTCTTCCGCCAGTACG | ACGATGGGTAATAGGCCGAG | 128 | 64.0 | | 63.1 | 60.0 | 55.0 | 33.0 | 11.7 | 11.6 | Designed |
| *TNFα* | NM_000594.4 | GCCCATGTTGTAGCAAACCC | TGAGGTACAGGCCCTCTGAT | 133 | 63.7 | | 64.2 | 55.0 | 55.0 | 30.6 | 13.4 | 15.4 | Designed |
| *TNFα* | NM_000594.4 | CCTCTCTGCCATCAAGAGCC | TCCCAAAGTAGACCTGCCCA | 182 | 60.2 | | 60.5 | 60.0 | 55.0 | 40.2 | 12.2 | 8.0 | Designed |
| *TNFα* | E00702.1 | GGCAGTCAGATCATCTTCTCG | ATGGCAGAGAGGAGGTTGAC | 294 | 58.3 | | 59.1 | 52.4 | 55.0 | 12.5 | 12.3 | 12.9 | (Edwards et al., 2000) |
| *TPPP* | NM_007030.3 | AGGACTGCCAGGTGATCGAC | CTTGCCCTCGATGAGCCTGT | 200 | 65.5 | | 66.1 | 60.0 | 60.0 | 40.1 | 17.6 | 20.2 | (Inokawa et al., 2016) |
| *TPPP* | NM_007030.3 | AAGAGGCTGTCGCTGGAATC | TGATGGTCCGGCAAGACTTC | 250 | 64.1 | | 64.1 | 55.0 | 55.0 | 38.2 | 9.2 | 11.8 | Designed |
