## Supplementary Table S5 for "The glia club: Validation of polarization biomarkers for human microglia (HMC3) using quantitative real time RT-qPCR"

**Supplementary Table S4.** Primer sequences and RT-qPCR parameters for candidate HMC3 reference genes with differential expression between experimental groups. Targets are identified by gene name, NCBI accession number, targeted transcript variants, forward and reverse primer sequences, annealing temperature (°C), cDNA amount (ng), threshold cycle (Ct), product size (bp), and function. The sources of the primers (designed using NCBI or obtained from literature) are indicated. All primers target *Homo sapiens*.

| **Gene** | **NCBI Accession #** | **Variants targeted** | **Forward primer (5’ – 3’)** | **Reverse primer (5’ – 3’)** | **Annealing temperature (℃)** | **cDNA loaded (ng)** | **Ct range*^a^*** | **Product size (bp)** | **Function** | **Source of primer** |
| --- | --- | --- | --- | --- | --- | --- | --- | --- | --- | --- |
| *ACT*$B$ | NM_001101.5 | N/A | GGAGTCCTGTGGCATCCACG | CTAGAAGCATTTGCGGTGGA | 63 | 10 | 18.3-19.4 | 322 | Regulates expression of genes involved in cell cycle and cell migration (Bunnell et al., 2011). | (Robertson et al., 1999) |
| *TPI1* | NM_001258026.2 | 1-3 | CGCAGATAACGTGAAGGAC | CAGTCACAGAGCCTCCATAA | 61 | 10 | 21.1-22.3 | 191 | Catalyzes the isomerization of dihydroxyacetone phosphate (DHAP) and glyceraldehyde-3-phosphate (GAP) in glycolysis (P. Liu et al., 2022). | (Hernández-Ochoa et al., 2021) |
| *PGK1* | NM_000291.4 | N/A | ATGGATGAGGTGGTGAAAGC | CAGTGCTCACATGGCTGACT | 63 | 10 | 23.3-24.7 | 118 | Catalyzes the reversible reaction of 1,3-bisphosphoglycerate (1,3-BPG) to 3-phosphoglycerate (3-PG) in glycolysis and gluconeogenesis (Bernstein & Hol, 1998; H. Liu et al., 2022). | (Carrillo-Jimenez et al., 2019) |
| *TKT1* | XM_054347704.1 | 1-3 | GATCACGGGGGTAGAAGA | TGTCCCCAACTTTGTAGCT | 61 | 10 | 23.5-24.0 | 200 | Catalyzes reactions in the non-oxidative branch of the pentose phosphate pathway (Alexander-Kaufman & Harper, 2009). | (Hernández-Ochoa et al., 2021) |

*a)* The indicated Ct range reflects the amplification of control HMC3 cells
